# ChromWave: Deciphering the DNA-encoded competition between transcription factors and nucleosomes with deep neural networks

**DOI:** 10.1101/2021.03.19.436198

**Authors:** Sera Aylin Cakiroglu, Sebastian Steinhauser, Jon Smith, Wei Xing, Nicholas M. Luscombe

## Abstract

Transcription factors (TFs) regulate gene expression by recognising and binding specific DNA sequences. At times, these regulatory elements may be occluded by nucleosomes, making them inaccessible for TF-binding. The competition for DNA occupancy between TFs and nucleosomes, and associated gene regulatory outputs, are important consequences of the cis-regulatory information encoded in the genome. However, these sequence patterns are subtle and remain difficult to interpret. Here, we introduce ChromWave, a deep-learning model that, for the first time, predicts the competing profiles for TF and nucleosomes occupancies with remarkable accuracy. Models trained using short- and long-fragment MNase-Seq data successfully learn the sequence preferences underlying TF and nucleosome occupancies across the entire yeast genome. They recapitulate nucleosome evictions from regions containing “strong” TF binding sites and knock-out simulations show nucleosomes gaining occupancy in the absence of these TFs, accompanied by lateral rearrangement of adjacent nucleosomes. At a local level, models anticipate with high accuracy the outcomes of detailed experimental analysis of partially unwrapped nucleosomes at the GAL4 UAS locus. Finally, we trained a ChromWave model that successfully predicts nucleosome positions at promoters in the human genome. We find that human promoters generally contain few sites at which simple sequence changes can alter nucleosome occupancies and that these positions align well with causal variants linked to DNase hypersensitivity.

## 1 Introduction

Transcription factors (TFs) and nucleosomes play critical roles in regulating gene expression. TFs recognise and bind short, specific DNA sequences and occupy thousands of discrete sites in the genome. Nucleosomes on the other hand - comprising a 147bp section of DNA wound around a histone octamer - cover most of the genome with a preference for GC-rich sequence regions. With some exceptions, binding of a stretch of DNA by TFs and nucleosomes is generally considered to be mutually exclusive (Hayes and Wolffe, 1992; Zhu *et al*., 2018) and histones are evicted or repositioned to expose or occlude TF-binding sites. Thus, the competition between TF and nucleosome occupancies plays an important role in effecting gene regulatory outcomes (Svaren *et al*., 1994; Almouzni and Wolffe, 1995; John *et al*., 2008, 2011; Kim and O’Shea, 2008; Lam, Steger and O’Shea, 2008; Mirny, 2009; Kaplan *et al*., 2011; Neph *et al*., 2012; Raveh-Sadka *et al*., 2012; He *et al*., 2013). However, genome-wide understanding of DNA sequence information that determines the competition between these two sets of molecules remains elusive.

Existing sequence-based computational models successfully predicted either TF-binding (e.g. via position weight matrices; (Wasserman and Sandelin, 2004; Mathelier and Wasserman, 2013; Jayaram, Usvyat and R Martin, 2016) or nucleosome occupancies individually (e.g. using Hidden Markov Models, HMM; (Segal *et al*., 2006; Peckham *et al*., 2007; Gupta *et al*., 2008; Kaplan *et al*., 2009). HMMs have also been applied to predict competitive multi-factor binding in yeast (Wasson and Hartemink, 2009; Ozonov and van Nimwegen, 2013; Zhong, Wasson and Hartemink, 2014) as well as logistic regression (He *et al*., 2013). However, key underlying assumptions of HMMs are violated, notably that (i) all DNA-binding proteins and their binding motifs are known, (ii) binding sites and probability of binding are fully described by the motif alone, and (iii) the binding strength for each protein and each motif is known. HMMs do not allow for easy integration of various data types measuring different influences on TF and nucleosome occupancies (e.g. DNA-sequence, 3D conformation of the genome, histone marks, DNA methylation). Moreover, the computational effort required to compute the right concentrations for even just a few TFs is substantial. Such drawbacks limit the accuracy and application of such models.

Recent advances in the application of deep learning in genomics have shown great promise in modelling several signals (often simultaneously) from DNA sequence alone (Alipanahi *et al*., 2015; Angermueller *et al*., 2017; Zeng and Gifford, 2017; Kelley *et al*., 2018; Wang *et al*., 2018; Avsec, Weilert, *et al*., 2021). These results make a unified model that can integrate all these different data to predict gene regulation genome-wide conceivable. However, most existing approaches score a whole DNA-sequence with the probability of a single binding event by one DNA-binding factor such as TFs, RNAs or nucleosomes and do not predict two cometing binding events at the same time along a sequence as previous HMM-based models. In addition, all except (Avsec, Weilert, *et al*., 2021) apply predictions to bins of 100 nucleotides or more, thus limiting their applications to interpreting the effects of local sequence changes such as single nucleotide polymorphisms.

Here, we present ChromWave, a flexible deep learning model to understand the effect of genetic variation on DNA binding at nucleotide-resolution. Our approach substantially improves the current gold standard of models and alleviates some of their constraining assumptions, such as the knowledge of all DNA-binding proteins together with their sequence specificity. At the expense of the ability to distinguish different TF binding events, ChromWave can predict nucleosome- and TF-binding genome-wide, and can resolve binding events not directly explainable by an annotated motif.

## 2 Results

### 2.1 The ChromWave deep-learning architecture

ChromWave is a deep-learning algorithm to predict competing TF and nucleosome-occupancies at nucleotide resolution on a genome-wide scale. It is based on Google Deepmind’s WaveNet model (van den Oord, Dieleman, *et al*., 2016) that interprets textual information from many adjacent positions to infer an audio output. The input to ChromWave is a genomic sequence and its outputs are one or more discretised binding profiles of the same length. The number and types of outputs can be varied (for example, TFs v nucleosomes, nucleosomes in different conditions). Predictions are conditioned on several kilobases of sequence spanning the region of interest, meaning that nucleotides surrounding a site are taken into account. Models are readily trained using data from experimental measurements of genomic occupancies (such as ChIP-seq, CUT&RUN, MNase-seq, or ATAC-seq): data can be presented as processed log-ratio values or raw count data with minimal preprocessing (Methods).

ChromWave accepts large sequence regions as input (Figure 1A; “Input”); in the example below, training was performed using 5 Kb sequences, and predictions were made across whole chromosomes. The model applies one layer of convolutions to transform the DNA sequence into a series of vectors of the same length (Figure 1B; “Convolution Layers”). This first layer of convolution learns patterns in the DNA sequence that can be considered to be motifs that are important for model predictions; we refer to these learned patterns as motif filters (Alipanahi *et al*., 2015). Transfer-learning can be applied at this stage - for instance, to incorporate models trained on another data set or on data from another organism - by combining the first convolutional layer with a pre-trained layer and concatenating their outputs.

**Figure 1.**
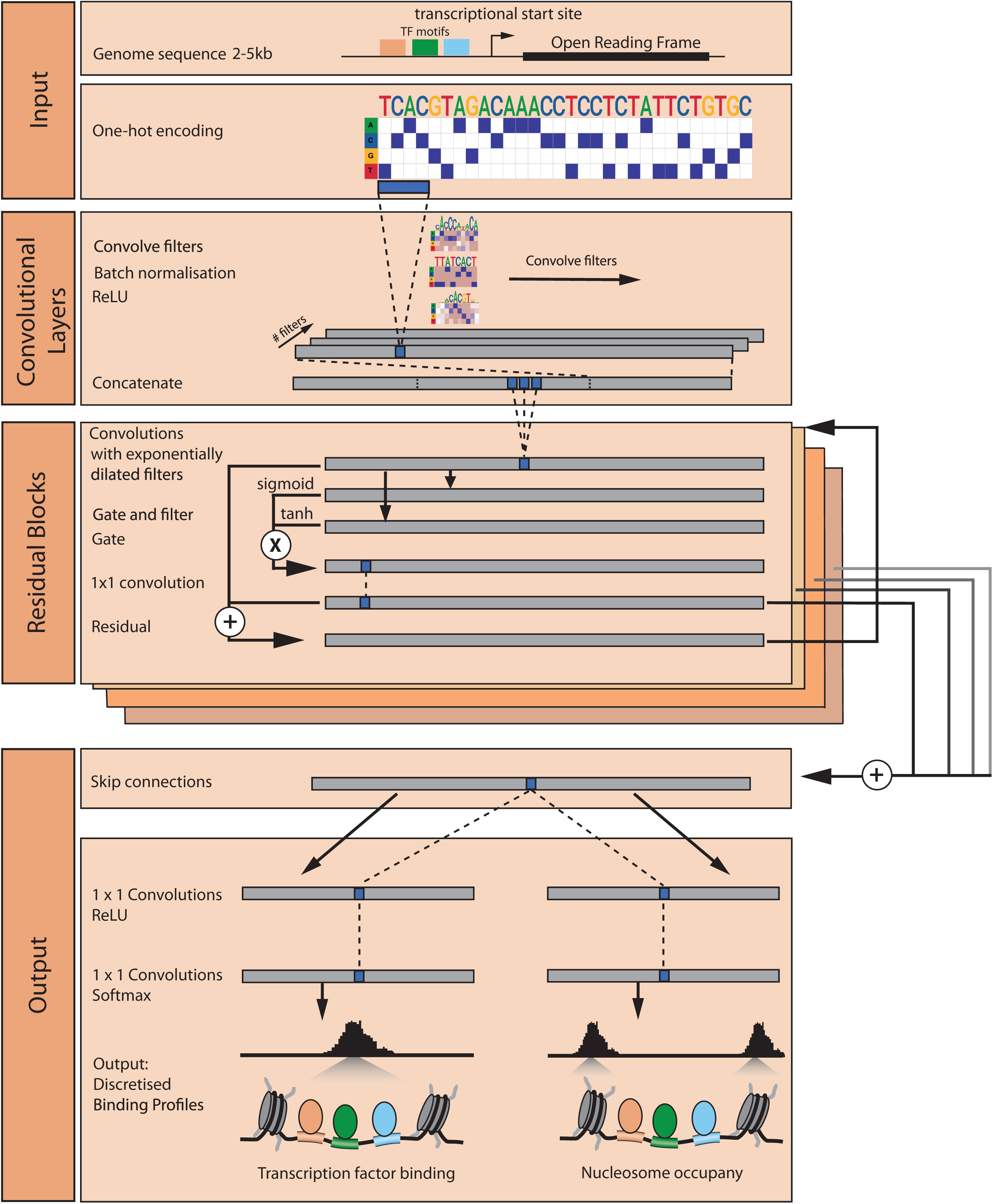
The ChromWave deep learning architecture to predict chromatin accessibility profiles. **Input.** DNA sequences are one-hot coded to four rows representing A, C, G, and T before entering the model. The annotations of the transcription factor binding sites and transcriptional start site are examples indicated to help convey the reasons for the various elements of the architecture. **Convolutional Layers.** We first apply a convolutional layer followed by batch normalisation and a ReLU layer. If we used transfer-learning we also use a convolutional layer with pre-trained weights followed by a max-pooling layer before concatenating these with the trainable convolutional layer. **Residual Blocks.** To share information across large distances, we apply several layers of dilated convolutions in at least nine residual blocks. Each residual block contains two dilated convolutions with exponentially dilating filters (one gate, one filter) followed by element-wise multiplication. The gated activation output is (i) outputted as skip connections which are aggregated after all residual blocks, and (ii) as residual summed with the block input and used as input to the next residual block. **Output.** After the final residual block and aggregating the skip connections, for each output profile a convolutional layer with kernel size 1, stride of 1, and ReLU activation is applied and passed to the final prediction layer where another convolution layer with kernel size 1, stride of 1 makes predictions across the sequence. The output therefore is of dimension (sequence length) x (number of bins).A prediction is made for each input position by the last layer through a softmax activation function applied to the second dimension and represents the predicted bin of the discretised binding profiles. We compare these predictions to the experimentally observed profiles (the target) via the mutual information between observed and predicted classes, as well as the Pearson correlation coefficient (the latter is not used during the fitting of the model parameters with stochastic gradient descent with back propagation, but only for the selection of the best model during the hyperparameter optimization).The predicted (discretised) profiles are finally turned into continuous profiles by reversing the discretisation process used on the input data.

Next, to share information across neighbouring regions, we apply multiple layers of dilated convolution by stacking residual blocks (Figure 1C; “Residual Blocks”); these can be considered the main building blocks of ChromWave and are similar to those of WaveNet (van den Oord, Dieleman, *et al*., 2016) and DeepC (Schwessinger *et al*., 2020). A residual block consists of two dilated convolutional layers with exponentially dilating filters, followed by element-wise multiplication. The first residual block accepts the concatenated output of the first convolutional layer(s) as input and subsequent residual blocks accept the output of the previous one as input. In this way, local features in lower dilated convolutional layers are accumulated sequentially to capture dependencies between distal sequence locations up- and downstream.

Each residual block produces two outputs: (i) a feature map from the previous residual block to be used as input to the next one and (ii) a skip connection that is used to calculate the loss function (for the input batch) once all the residual blocks have been processed. Within a residual block, the input passes through a gated activation z=sigmoid(x) ⊙ tanh(x), where ⊙ denotes the element-wise multiplication (similar to the LSTM gated activation units used in PixelCNN; (van den Oord, Kalchbrenner, *et al*., 2016). Similar to the WaveNet architecture (van den Oord, Dieleman, *et al*., 2016), the gated activation output is finally summed element-wise with the block-input to form the residual. Residuals and skip connections allow faster convergence and stable training for deeper networks (He *et al*., 2016).

Finally, for each output profile, we apply a width-one convolutional layer with a Rectified Linear Unit (ReLU) activation (Figure 1D; “Output”). ChromWave outputs a categorical distribution over the discretized binding profile values with a softmax layer, optimized to maximize the negative log-likelihood (for example, multi-class cross-entropy) of the data with respect to the parameters. In the case of multiple output profiles, the loss function is the sum of the individual cross-entropies for each profile. We tune hyperparameters on a validation set and we can easily gauge if the model is over- or underfitting, while we use a held-out test set for final validation after training.

For each nucleotide position, ChromWave predicts an occupancy class that represents the binned read count. However, the distribution of bins tends to be highly imbalanced, with most representing the genomic average. An advantage of using classification over regression is that we can weight correct or false predictions of rarer classes with very high or low read counts to deter the model from defaulting to the most frequent bins. We achieve this by applying a weighted multi-class cross-entropy using the median/frequency per class as weight (Methods). Finally, we map ChromWave’s predictions of the discretized binned read counts back to the original continuous value ranges (which we generally refer to as ChromWave’s predictions unless explicitly otherwise stated).

### 2.2 ChromWave performs a nucleotide-resolution prediction of TF and nucleosome occupancies across the whole yeast genome

We trained a model of TF and nucleosome occupancies for the yeast genome using paired-end MNase-seq data from (Henikoff *et al*., 2011). The dataset contains multiple fragment lengths (20-200bp) reflecting the diverse footprint sizes of the bound proteins and protein complexes. We separated fragments into long (140–200 bp) and short (20-80bp) groups as in (Zhong, Wasson and Hartemink, 2014). Note that while the short fragments are unlikely to represent unwrapped nucleosomes (Henikoff *et al*., 2011), they do not distinguish between TF types and could represent other small DNA-binding protein or protein complexes that are not TFs. However, for the ease of notation, we will denote these as nucleosome and TF occupancies respectively. We used the numbers of read fragments that cover each nucleotide position as the measure of nucleosome and TF occupancies (Zhong, Wasson and Hartemink, 2014).

We provided numerous alternative transfer-learning strategies as options to our hyperparameter search (Methods). One of these options was to include 256 pre-trained convolutional filters with kernel size 6. These filters were pre-trained as part of a binary classification convolutional neural network to predict nucleosome dyads mapped by chemical cleavage (Brogaard *et al*., 2012). The best ChromWave model includes these 256 pre-trained convolutional filters and another 256 trainable convolutional filters with kernel size 16. Another option to our hyperparameter search that was chosen by the best solution was to initialize 56 of these convolutional filters with position weight matrices of 28 known nucleosome-displacing transcription factors and their reverse complements: Abf1, Cbf1, McM1, Rap1, Reb1, Orc1, Asg1, Azf1, Bas1, Ecm22, Ino4, Leu3, Rfx1, Rgm1, Rgt1, Rsc3, Sfp1, Stb4, Stb5, Stp1, Sum1, Sut1, Tbf1, Tbs1, Tea1, Uga3, Ume6, and Urc2 (Yan, Chen and Bai, 2018).

Figure 2A provides the first view of ChromWave’s prediction in three example regions. The model has produced two output profiles: one for nucleosome occupancies and another for TF-binding. The first two examples illustrate the good agreement of the predicted occupancies in black, with the original MNase-seq signal in grey. nucleosome occupancies follow the well-known periodic profile, with the typical interruptions found in the nucleosome-depleted regions of promoters (red stippled boxes). Interspersed between the breaks in nucleosomes, it is possible to observe TF-binding. We examine the competition between nucleosomes and TFs below. The third example region shows a region lacking MNase-seq reads in the original data, e.g. were not covered by any high-quality alignments. The most common reason for this effect often seen across various NGS datasets is high repetitiveness in the region so that no read can be attributed to any specific location with high confidence. ChromWave imputes both TFs and nucleosomes with plausible binding profiles in such cases.

**Figure 2.**
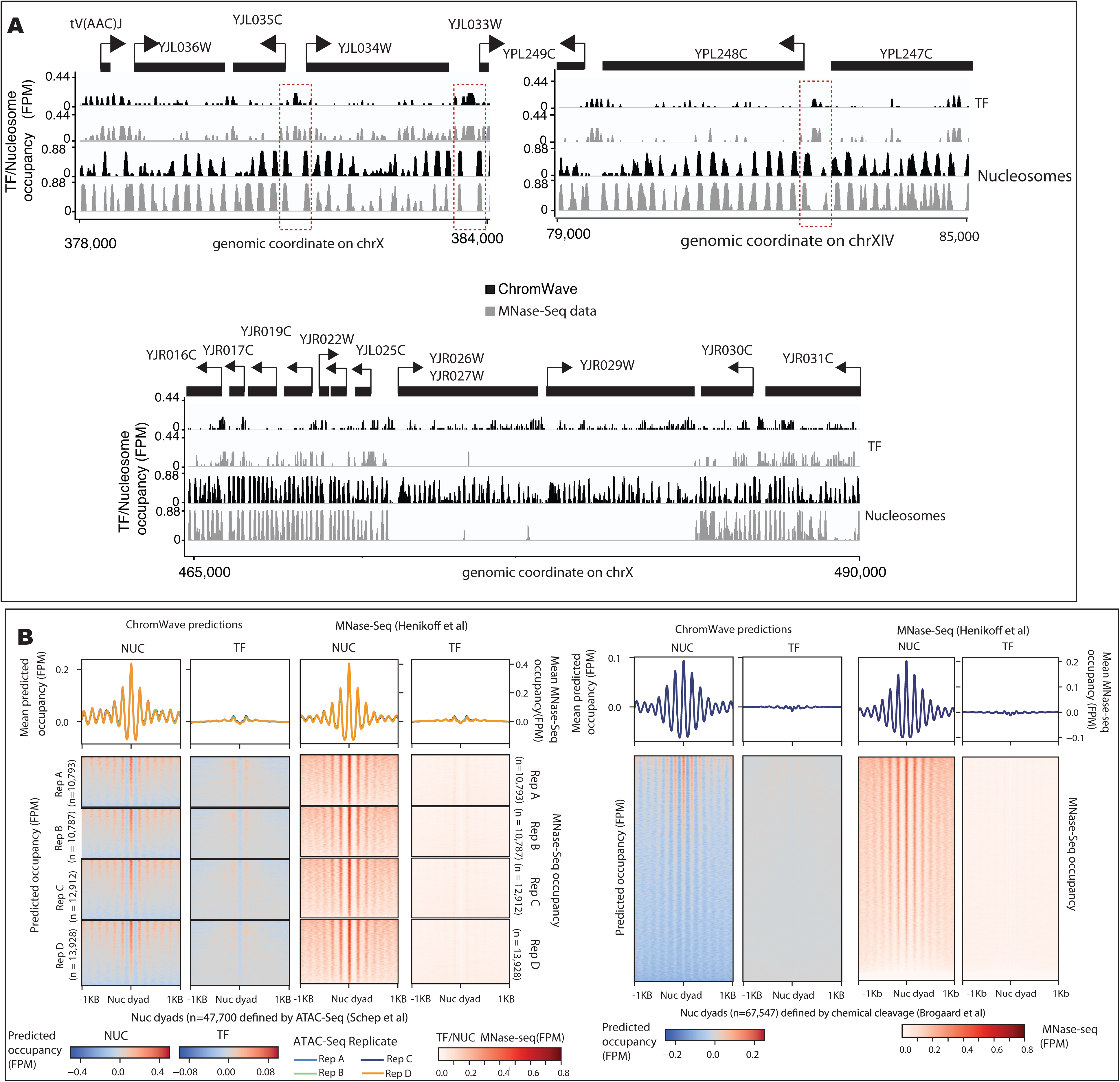
Nucleotide-resolution prediction of TF and nucleosome occupancies in the yeast genome. **A.** Top: Two representative examples of the predicted TF and nucleosome occupancy profiles from the training set (chromosome X) and the unseen test set (chromosome XVI). The profiles predicted by ChromWave are shown in black for comparison with the observed MNase-seq data shown in grey. Bottom: A representative example imputed TF and nucleosome occupancies from the training set (chromosome X). Shown are ChromWave’s predictions of TF and nucleosome occupancy profiles (in FPM) in black in an unmapped region for comparison with the observed MNase-seq data (in FPM) shown in grey. **B.** ChromWave’s predictions of TF and nucleosome occupancy around experimentally defined nucleosome dydas. Heatmaps showing +/−1kb regions around nucleosome dyads called either on ATAC-seq (left, (Schep *et al*., 2015)) or defined by chemical cleavage (right, (Brogaard *et al*., 2012)) data. The ATAC-seq heatmap was split according to four biological replicates (A, B, C, D). Nucleosome occupancy (NUC) and TF-binding (TF) in FPM from ChromWave’s predictions and the MNase-Seq data are shown as separate heatmaps next to each other as indicated. Average predictions are shown as metaprofiles on top of each heatmap. Each track was normalised by subtracting the genomic average before plotting.

To assess the model’s predictions at nucleotide resolution, we tested ChromWave’s predictions against genome-wide *in vivo* nucleosome dyads identified with high precision using an alternative experimental approach (e.g. chemical cleavage) (Brogaard *et al*., 2012) ChromWave correctly predicts nucleosome occupancy compared with the derived dyads, including the phasing of nucleosomes (Figure 2B, left). To measure the model’s performance across the entire yeast genome, we mapped the predicted occupancy classes to the continuous value ranges of the original binned read counts and computed the geometric mean of the Pearson’s correlation coefficients between the predicted occupancies and the smoothed MNase-seq read counts for nucleosomes and TFs (Figures 3A and S1). It is immediately apparent that ChromWave achieves a good fit across all chromosomes in both the TF and nucleosome profiles (genome-wide Pearson correlation of 0.65 and 0.49 for TFs and nucleosomes respectively) with no apparent biases towards specific chromosomal regions such as centromeres and telomeres (Figures 3A and S1B). Black regions correspond to those lacking MNase-seq reads and thus comparisons could not be made.

**Figure 3.**
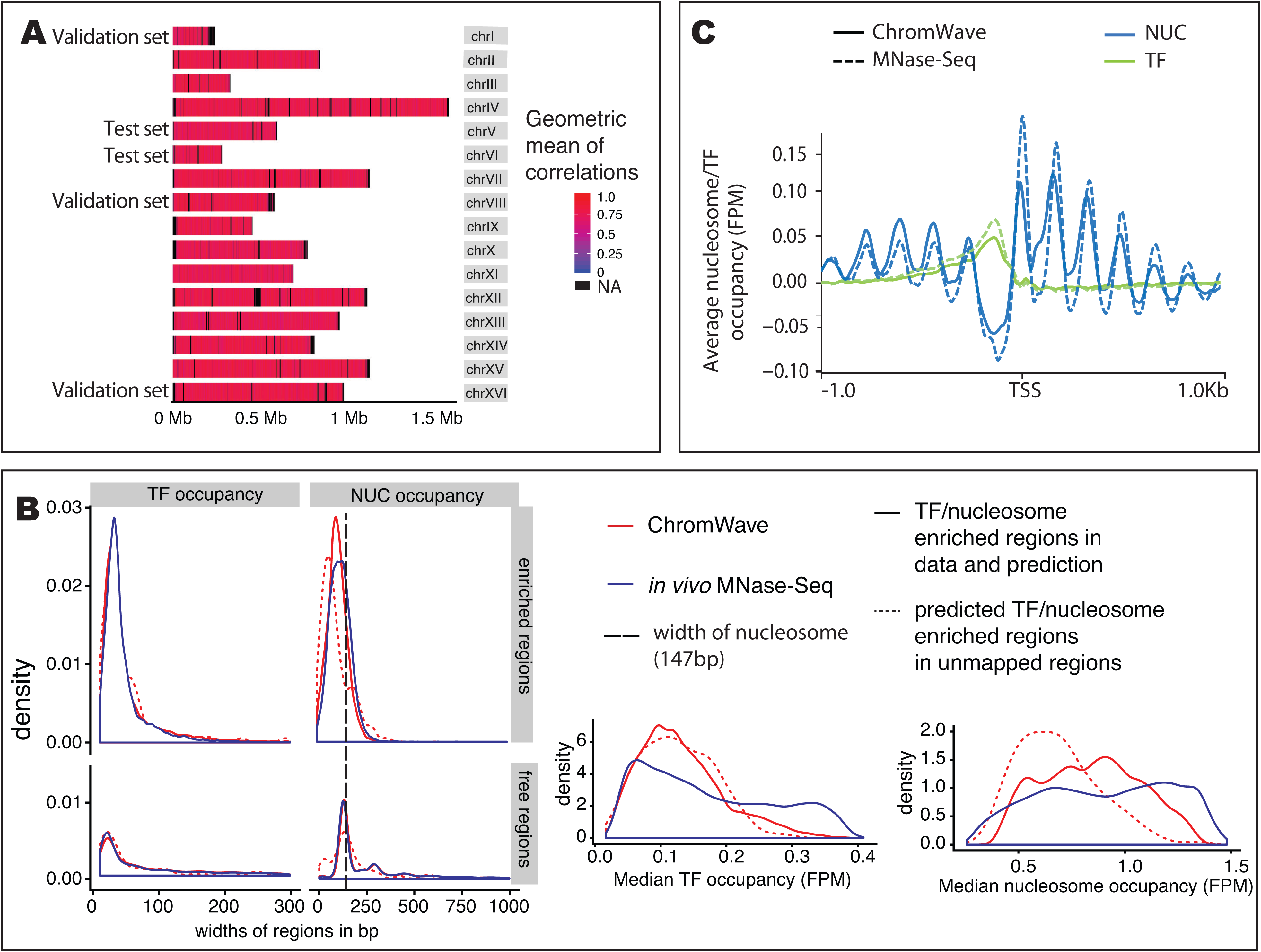
ChromWave’s accurately predicts nucleosome occupancy and TF-binding in mapped and unmapped regions. **A.** Geometric means of the Pearson Correlation coefficients between predicted and observed TF and nucleosome occupancy genome-wide. Indicated are which chromosomes formed the validation and test sets (all other chromosomes formed the training set). Correlations were computed in sliding windows of size 300bp and then averaged over 300bp bins. **B.** Left: Density plots of lengths of bound (eg ‘enriched regions’) and unbound regions (eg ‘free regions’) for TF and nucleosome predictions from data and predictions in mapped and unmapped regions. Right: Density plots of median signal amplitudes for TF and nucleosomes (called as ‘enriched’ regions) from data and predictions in mapped and unmapped regions. To plot signal amplitudes we applied a jitter of 0.4 to ChromWave’s predictions to smooth out the discrete scale for visualisation. **C.** Average predicted and observed TF and nucleosome occupancy in yeast promoters. Metaprofiles +/− 1KB of ChromWave’s predictions and MNase-seq data (in Fragments Per Million, FPM) around all TSSs for predicted and observed TF and nucleosome occupancy. Each track was normalised by subtracting the genomic average before plotting.

In terms of the discretized occupancy classes, ChromWave tends to predict with higher precision classes corresponding to higher read count bins and there is a general tendency to overestimate occupancies (Figure S1A). To assess prediction accuracies of the locations of binding sites and nucleosome positions, we defined ‘enriched’ regions of 10 bp and 50bp where the average signal exceeded the 85th percentile for the TF and nucleosome profiles respectively. We compared the enriched regions thus defined in both predicted and observed profiles and found high accuracies in terms of positions (Area-under-the-ROC-Curve when predicting enriched regions were 0.75 for both TF and nucleosome predictions). Signal amplitude and spacing (Figure 3B) was consistent between data and predicted profiles. The results also suggest that the amplitude and spacing profiles of predictions in the unmapped regions are realistic.

### 2.3 ChromWave successfully predicts the competition in DNA occupancies between TFs and nucleosomes

Figures 2A and 3A provide encouraging signs that ChromWave is able to model the competition for DNA-binding between nucleosomes and TFs. The genome-browser tracks show several nucleosome-free regions (5’ NFRs) just upstream of transcriptional start sites (TSSs; indicated by stippled boxes). These regions are known distinguishing features between *in vitro* and *in vivo* nucleosome occupancy maps that arise through the binding of TFs and transcription initiation factors.

To assess whether ChromWave has learned a general promoter architecture rather than an implicit competition between TFs and nucleosomes, we further tested ChromWave’s predictions against genome-wide *in vivo* nucleosome dyads identified using alternative experimental approaches, chemical cleavage (Brogaard *et al*., 2012) and ATAC-Seq (Schep *et al*., 2015). Whereas ATAC-Seq lacks coverage in heterochromatin (and thus, defined dyads are located in accessible chromatin only), chemical cleavage maps dyads genome-wide. ChromWave correctly predicts nucleosome occupancy compared with the derived dyads from ATAC-Seq, including the phasing of nucleosomes (Figure 2B). Notably, there is a depletion of TF signal directly on the dyad in both prediction sets, indicating that ChromWave has learned that either TFs or nucleosomes can be bound at a DNA site at a time. At a global level, the meta-profiles of TF and nucleosome occupancies demonstrate that ChromWave accurately captures profiles around TSSs (Figure 3C).

### 2.4 ChromWave uses regulatory motifs to model the competition between TFs and nucleosomes

A class of abundant and essential TFs including Abf1, Rap1, Reb1, and Cbf1 drive promoter nucleosome exclusion around TSSs (Kent *et al*., 1994; Yarragudi *et al*., 2004; Badis *et al*., 2008; Hartley and Madhani, 2009; Tsankov *et al*., 2010, 2011; Ganapathi *et al*., 2011). In order to assess whether ChromWave successfully models the competition between TFs and nucleosomes, we examined ChromWave’s first convolutional layer to search for learned DNA sequence patterns that are important in predicting nucleosome and TF occupancies.

The short fragments in the input MNase-seq data do not distinguish between TF types, therefore, we inspected the learned sequence motifs to see if we could match the predicted occupancy positions to specific TFs. We examined the first convolutional layers of ChromWave that take the one-hot encoded DNA sequences as their input. Each filter can be summarised as a conventional position frequency matrix and visualised as a seqlogo plot as previously described (Alipanahi *et al*., 2015; Angermueller *et al*., 2017) searched these matrices against known yeast TF motifs (Zhu and Zhang, 1999; MacIsaac *et al*., 2006; Gupta *et al*., 2007; Pachkov *et al*., 2013; Hume *et al*., 2015; Fornes *et al*., 2020). In total, 36 of 256 filters matched known motifs (q-value < 0.05) including only 6 the 56 filters that were intialised TF motifs and their reverse complements. However, these initialised filters did not match their original motif, and instead were like the remaining filters augmented during training to capture information that is useful for the prediction task. Among the learned sequences are those that match motifs for Abf1, Rap1, and Reb1 (Figure 4A; Cbf1 did not have a good match).

**Figure 4.**
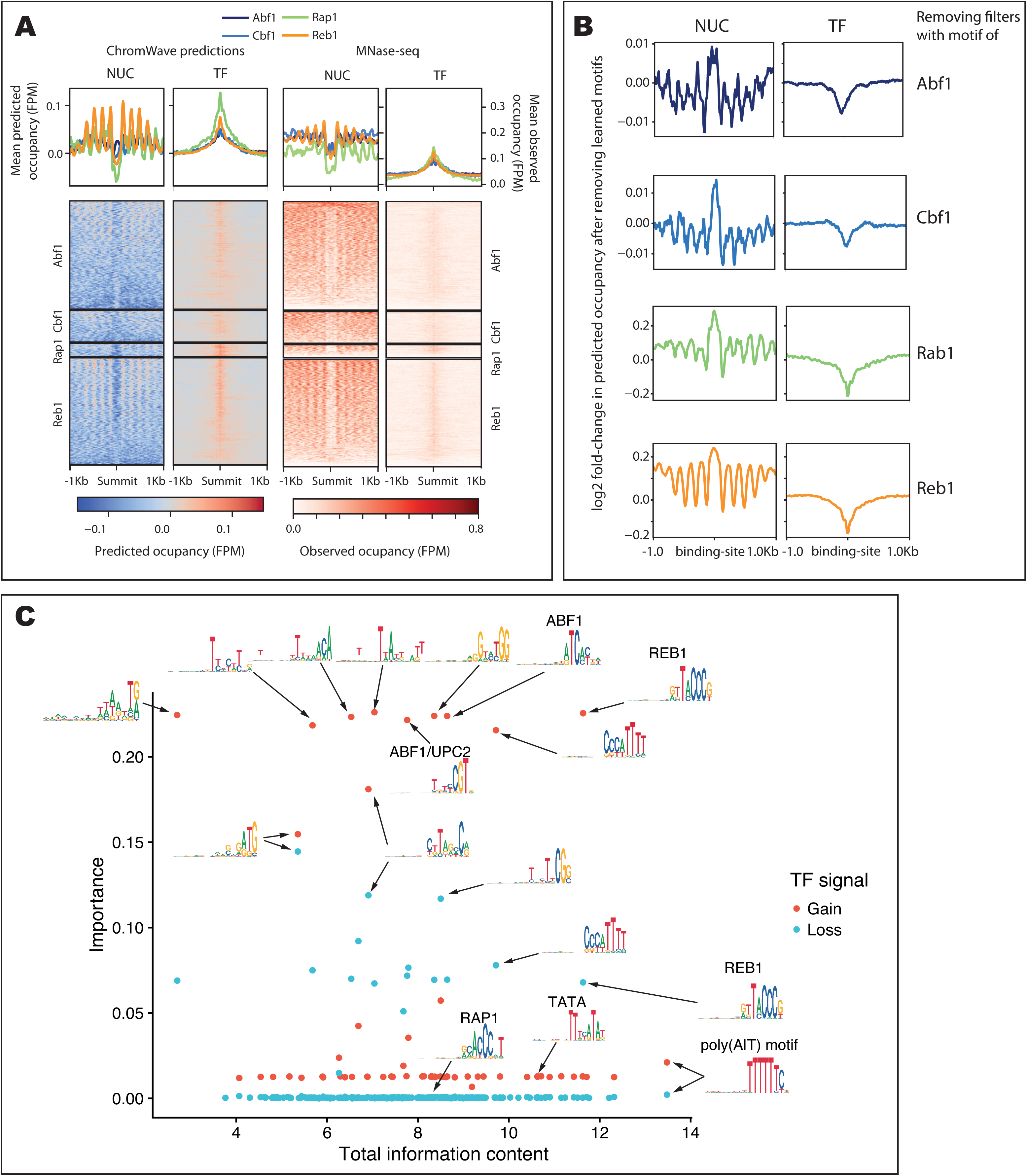
ChromWave uses regulatory motifs to model the competition between transcription factors, chromatin remodelers and nucleosomes. **A.** Nucleosome and TF occupancy per base-pair centered on TF binding sites defined by ChIP-exo. Heatmaps showing +/−1kb regions around ChIP-exo peak. Each heatmap panel is based on a different TF peak set from top to bottom: Abf1, Cbf1, Rap1 and Reb1. Average predictions are shown as metaprofiles on top of each heatmap. Heatmaps showing ChromWave’s predictions (*left)* and MNase-seq data (*right)* of nucleosome (*NUC*) and TF (*TF*) occupancy as indicated (in Fragments Per Million, FPM). Each prediction track was normalised by subtracting the genomic average before plotting. **B.** Log2-fold change in nucleosome and TF occupancy predictions (in FPM) when ChromWave’s fillters are perturbed per base-pair centered on TF binding sites defined by ChIP-exo. Meta-profiles showing average log2-fold changes in +/−1kb regions around ChIP-exo peak. Each panel is based on a different TF peak set from top to bottom: Abf1, Cbf1, Rap1 and Reb1. Log2-fold change was computed between the predictions of the full ChromWave model and a perturbed model where the convolutional filters corresponding to the respective TF-binding motifs (defined by TOMTOM annotation) were set to zero.C. Importance of motif detectors associated with nucleosome-TF competition. Motif importance (setting associated filter to zero) was plotted against the sum of information content (IC) for each motif detector. The color indicates a gain or loss of TF binding prediction by setting the motif to zero. Selected motifs were annotated with their associated sequence logo.

To assess ChromWave’s abilities to model the competition between nucleosomes and specific TFs, we extracted previously published binding sites for Abf1, Rap1, Reb1, and Cbf1 derived from ChIP-exo assays (Rossi, Lai and Pugh, 2018) and aligned ChromWave’s outputs (Figure 4A). ChromWave predicts remarkably accurately nucleosome exclusion and TF occupancies at these sites with the largest effects seen for Rap1 and Reb1(Figure 4A). To understand how ChromWave relies on the learned TF motifs for its predictions, we set the weights of filters that had learned the motifs for Abf1, Rap1, Reb1, or Cbf1 individually to zero to mimic knock-out mutants. Figure 4B displays the difference in predictions between the mutant and wild-type conditions. For all four TFs, ChromWave predicts a loss of TF-occupancies and a corresponding increase of nucleosome occupancies. Taken together, these results indicate that ChromWave has learned that the competition between nucleosomes and TFs is encoded in the underlying DNA sequence.

### 2.5 ChromWave learns known and unknown regulatory motifs associated with DNA occupancy competition in S.cerevisiae

Next, we assessed the other learned sequences (Figure 4C). To derive a measure of importance, we set each filter matching a TF motif to zero and computed the absolute difference of the predicted TF occupancies compared with the full model at each bp position across the genome. To summarise all signal changes we obtain the value for the maximal increase (‘gains’) and decrease (‘loss’) in predicted TF occupancies within 300bp windows across the genome. To derive a single score for each TF, we compute the median of the maximal increase and decreases across all windows. A high importance value indicates a large contribution to predictions: a ‘loss’ value corresponds to a loss of a predicted TF peak upon deletion of the filter, whereas a ‘gain’ represents an increase in predicted occupancy.

ChromWave has learned some redundant filters, thus the removal of one filter may not perturb the model as it can be mitigated by other filters and would not be identified with this analysis. However, the removal of many filters changed ChromWave’s prediction dramatically: many of the matched and high-scoring motifs correspond to those of sequence-specific TFs that regulate nucleosome occupancy in 5’ NFRs such as Reb1 and Abf1 discussed above, as well as poly-(d:A,d:T) stretches (Figure 4C). These were also identified as some of the best-learned motifs of the convolutional layer (measured as the information content in the derived motif) together with several unknown motifs capturing lower-order sequence composition such as GC content. We also found additional motifs associated with TF binding that were deemed unmatched including several degenerated versions of the Reb1, the Rap1 motif, poly-(d:A,d:T) stretches, and TATA-box motifs.

### 2.6 ChromWave computationally recapitulates an experimental study of an unwrapped nucleosome between the Gal1 and Gal10 genes

Given these encouraging genome-wide results, we set out to investigate ChromWave’s performance at a detailed resolution on remodelled nucleosomes. One of the most abundant remodelers in yeast is Remodeling the Structure of Chromatin (RSC). In the Henikoff training data, an RSC-nucleosome complex is identified by shorter reads than for nucleosomes alone, as the DNA is unwound and becomes more susceptible to MNase digestion. Predicted occupancies for these remodelled nucleosomes thus are likely to be present in our TF profiles rather than the nucleosome profiles.

We focused on the GAL4 UAS in the GAL1-GAL10 region described as containing a partially unwrapped nucleosome which is remodelled by RSC (Floer *et al*., 2010). This nucleosome is apparent in the Henikoff MNase-seq dataset obtained at shorter digestion times (Henikoff *et al*., 2011) (Figure S1C). This peak has nearly disappeared in the 20-min digestion sample which we used for training ChromWave, whereas a peak presumably representing RSC or the RSC-nucleosome complex appears in the TF profile. As expected, ChromWave’s predictions on the yeast regulatory locus GAL4 UAS follow closely the training data: it predicts a broad TF peak representing the RSC-nucleosome complex directly in the UAS with strongly positioned nucleosomes on either side (Figure 5A).

**Figure 5.**
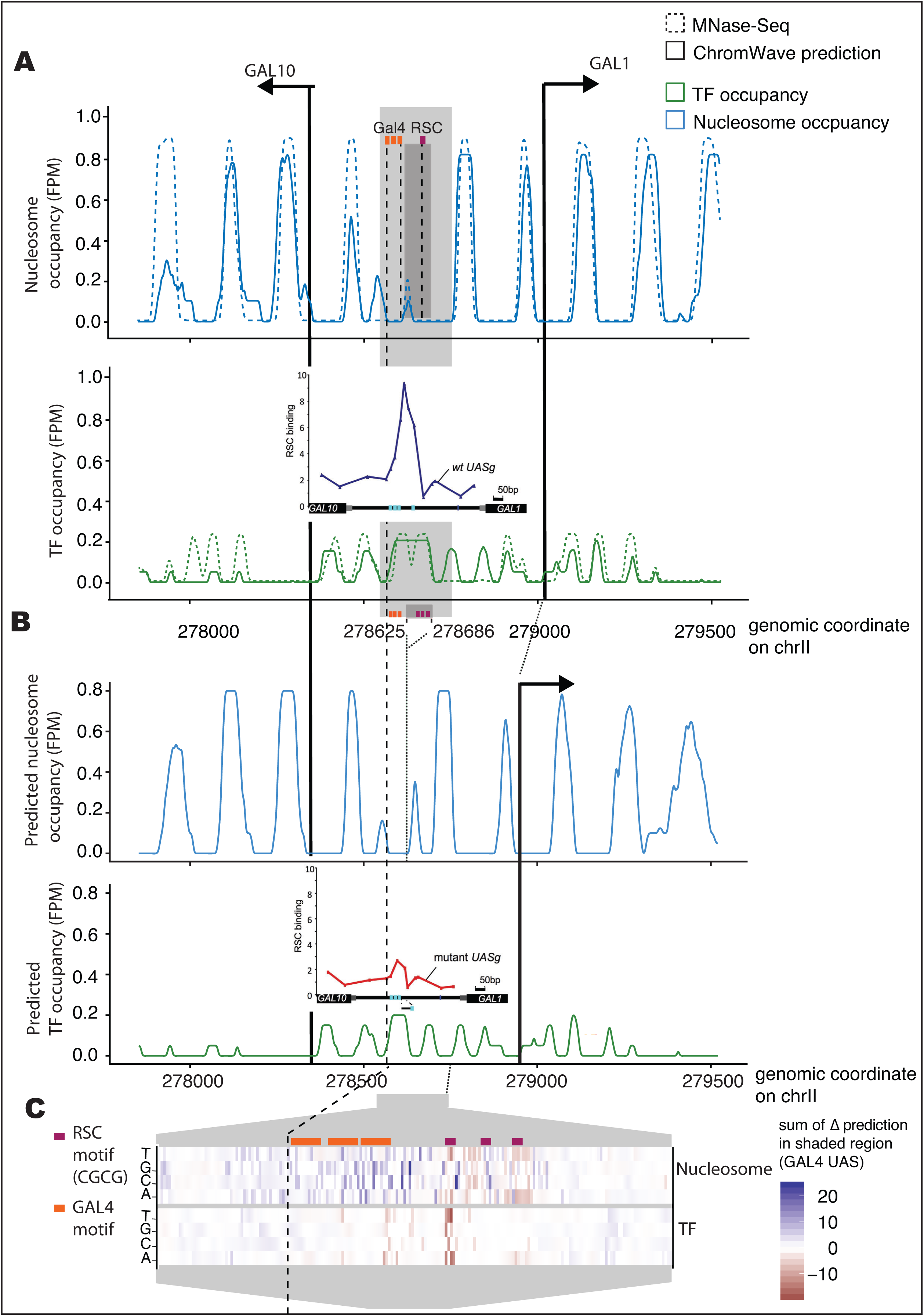
ChromWave recovers computationally an experimental study of an unwrapped nucleosome bound by RSC between the Gal1 and Gal10 genes. **A.** MNase-seq profiles and ChromWave’s predictions (in Fragments Per Million, FPM) of TF (green) and nucleosome (blue) occupancy in the intergenic region on chromosome II between the GAL10 and GAL1 genes on chromosome II. Annotations of Gal4 and RSC motifs are indicated by stippled lines. The light shaded region indicates a yeast regulatory locus GAL4 UAS. The dark shaded region indicates a 61bp region deleted inB. Inside, a partially unwrapped nucleosome bound by RSC is shown as a broad peak in the TF profile as the nucleosome is represented by shorter reads. *Inset:* Original Figure 1F from (Floer *et al*., 2010) for comparison. RSC binding to a WT *UASg*. Cells bearing TAP-tagged RSC and grown in raffinose. The four Gal4 sites within the *UASg* are in cyan. **B.** ChromWave’s predictions of TF (green) and nucleosome (blue) occupancy after deleting a 61bp sequence containing the RSC binding site (indicated by the stippled lines and the dark shaded region in A) in the GAL4 UAS. TF prediction changes to a narrow peak towards the 5’ end of GAL4 UAS, and a strongly positioned nucleosome is predicted in the middle of the GAL4 UAS. Previously hypersensitive sites on the borders of the shaded region are now predicted to be occluded by nucleosomes. These predictions recover experimental results in the original study on this unwrapped nucleosome (Floer *et al*., 2010). Inset: Original Figure 1F from (Floer et al., 2010) for comparison. RSC binding to the truncated UASg. Cells bearing TAP-tagged RSC and grown in raffinose. The four Gal4 sites within the UASg are in cyan. C. *In silico* mutagenesis with respect to the prediction in the Gal4 UAS (shaded region). Each column of the heatmap corresponds to a position in the sequence, each row represents a mutation to the corresponding nucleotide. The quantities in the heatmap display the difference “Δ pred” in ChromWave’s predictions (summed across the shaded region) after substituting the row’s specified nucleotide into the sequence. Annotation of the GAL4 and RSC motifs show that point mutations in a RSC binding site (which coincides with the first RSC motif) lead to the biggest loss in prediction in the TF occupancy profile in the GAL4 UAS.

The locus encodes three Gal4-binding sites as well as three putative binding sites of the RSC complex (Floer *et al*., 2010). We simulated the deletion of a 61bp sequence containing RSC3 binding motif performed by Floer (Floer *et al*., 2010). ChromWave correctly predicts a loss of TF occupancy at the deleted site and introduces a new narrower peak covering the three GAL4 motifs (Figure 5B). There is a dramatic increase in a narrow and strongly positioned nucleosome on the UAS, as well as nucleosomes encroaching from either side of the UAS. These nucleosomes now cover hypersensitive sites in wild-type that were reported to be increasingly occluded by nucleosomes upon RSC inactivation (Floer et al., 2010). Comparison with the molecular measurements by Floer highlights the remarkable accuracy of the ChromWave predictions (compare Figure 5B inset showing results from (Floer et al., 2010)).

To quantify how the underlying motifs and the surrounding sequence influences ChromWave’s prediction, we examined the changes in the prediction that arose upon perturbing the input sequence through *in silico* mutagenesis. At every nucleotide position in the GAL4 UAS, we considered mutations to each of the three other bases and computed changes in the predicted TF and nucleosome occupancies. For each position and each possible base, we then summed across all changes in the GAL4 UAS (Figure 5C). The largest effect is seen in a section region harbouring the 5’ most Rsc3 motif indicated in Figure 5C. Mutating the bases of the Rsc3 consensus motif contributed to the most extreme prediction changes in the TF profile: targeted disruption of the RSC motif by reducing the GC-content of the locus decreased the TF prediction in the shaded region dramatically. This putative binding site is also where RSC shows its maximal cross-linked ChIP signal (Figure 5A, Inset) (Floer *et al*., 2010).

### 2.7 Using ChromWave to quantify sequence patterns that differentiate between *in vitro* and *in vivo* nucleosome maps

Generally, only the nucleosome-sized MNase-seq fragments are retained in publicly available datasets and it will not always be possible to train a ChromWave model on both TF and nucleosome profiles. This is largely due to these data being more widely used to study nucleosome positioning as well as shorter fragments less easily interpretable since they can arise from many different proteins and protein complexes shielding the DNA from digestion by MNase. Therefore we set out to assess the predictions that can be made using models trained on different MNase-seq datasets.

Kaplan et al. published *in vitro* and *in vivo* nucleosome maps for the yeast genome (Kaplan *et al*., 2009), the former gauging the effects of DNA sequence alone on nucleosome-positioning and the latter including the impact of trans-acting factors. In the accompanying data of (Kaplan *et al*., 2009), nucleosome occupancy was measured as the log-ratio between the number of MNase-seq reads (of length ~ 147 bp) at a nucleotide position and the genome-wide average. In the same study, a Hidden Markov Model (HMM) trained on *in vitro* data was shown to predict *in vivo* nucleosome organisation with good accuracy, albeit with differences at transcription start sites (TSSs).

We first trained an *in vitro* ChromWave model using the *in vitro* MNase-seq dataset. We then trained an *in vivo* ChromWave model using the *in vivo* dataset but applying the pre-trained convolutional filters of the *in vitro* model as fixed filters (i.e. non-trainable), together with a further 256 trained convolutional layers. This approach gives us a direct way to detect differences in the learned sequence patterns between the *in vitro* and *in vivo* models.

Figure 6A shows examples of predictions from ChromWave’s *in vitro* and *in vivo* models and the Kaplan HMM (Kaplan *et al*., 2009), each producing a single output profile representing nucleosome occupancies. Overall, there is good agreement between all three model predictions and the MNase-seq data, with expected differences between the *in vivo* and *in vitro* profiles around TSSs (stippled boxes). At discretized occupancy class level, ChromWave’s *in vitro* and *in vivo* models show high prediction accuracy and precision (Figures S2A). Figure 6B gives an overview of ChromWave’s predictions across the entire genome when predicted classes representing read count bins are mapped back to the original continuous ranges: the predictions show good agreement with the observed data across all chromosomes with no apparent biases towards specific regions such as centromeres and telomeres. The lower correlation of the *in vitro* prediction on chromosome X is likely caused by artifacts in the original MNase-seq data (Figure S2B) and has been reported previously for other nucleosome positions models (Zhang *et al*., 2009; Liu *et al*., 2016). The average Pearson correlation with observed data genome-wide was 0.9 and 0.89 for *in vitro*- and *in vivo* nucleosomes respectively with a slightly increased performance in the training data for *in vivo* nucleosomes (Figure S2C). The average Pearson correlation genome-wide for the Kaplan HMM were 0.84 and 0.7 for *in vitro*- and *in vivo* nucleosomes respectively. We also compared the correlation of the predicted profiles with independently generated *in vivo* ATAC-seq datasets (Schep *et al*., 2015). Both ChromWave models perform better than the HMM, and the *in vivo* ChromWave model is far superior to the *in vitro* models (Figure S2C).

**Figure 6.**
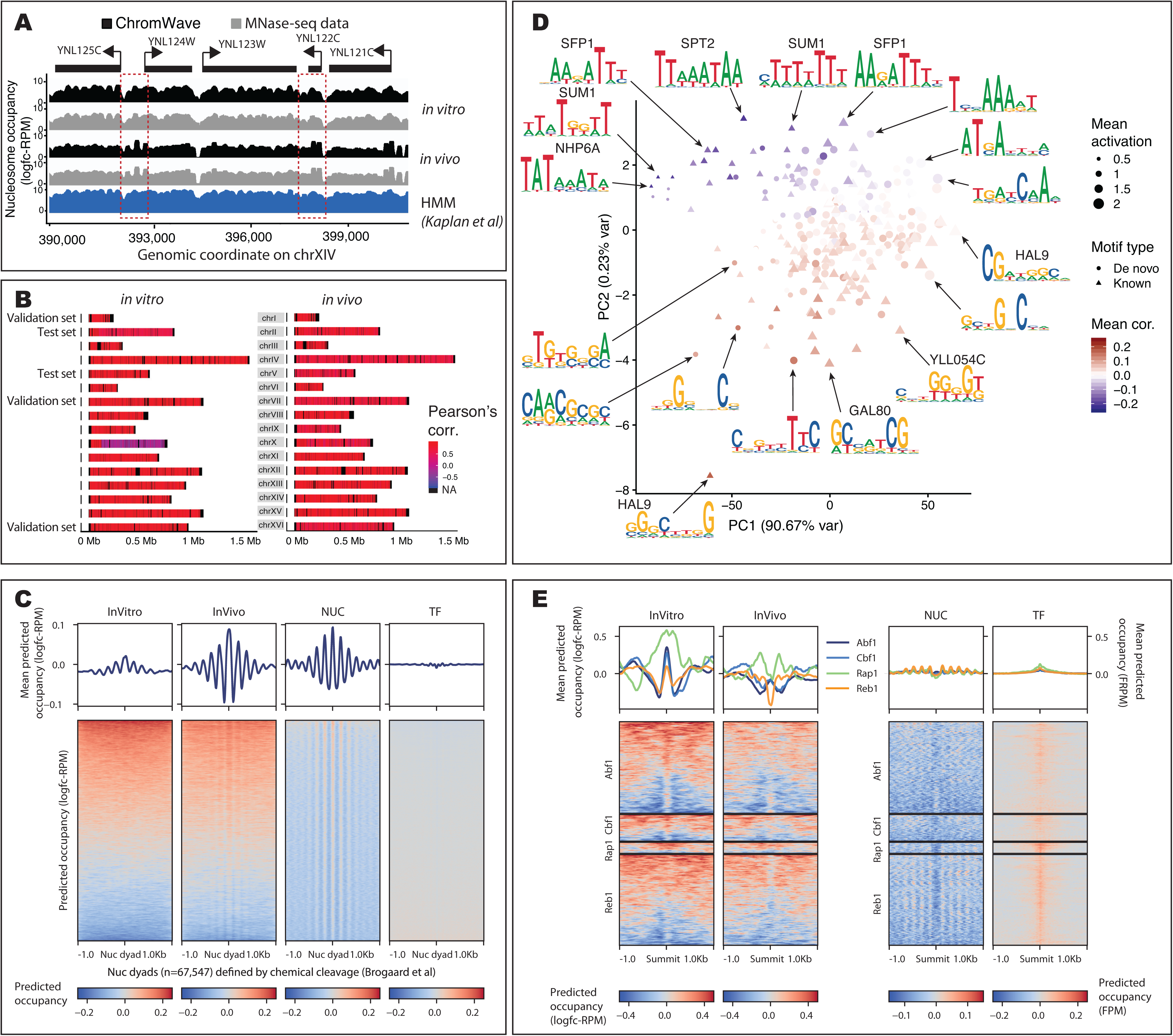
Nucleosome-only ChromWave models to decipher determinants of *in vivo* nucleosome occupancy in the yeast genome. **A.** A representative example of the predicted *in vitro* and *in vivo* nucleosome occupancy profiles from the training set (chromosome XIV). Occupancy is measured and predicted as log2-fold change of number of reads at each position over genome average, in Reads Per Million (logfc-RPM). Predictions by ChromWave are shown in black for comparison with the observed MNase-seq data shown in grey and the predictions of the Hidden Markov Model (HMM) trained on the *in vitro* data (Kaplan *et al*., 2009) shown in blue. Stippled boxes indicate observable differences in nucleosome occupancy *in vitro* and *in vivo.* **B** Genome-wide comparison of predicted and observed nucleosome occupancy. Pearson Correlation coefficients were computed between predicted and observed nucleosome occupancy (left: *in vitro,* right: *in vivo*). The chromosomes forming the validation and test sets as indicated (all other chromosomes formed the training set). Correlations were computed in sliding windows of size 300bp and then averaged over 300bp bins for visualisation. **C.** Predictions per base-pair centered on nucleosome dyads defined by chemical cleavage (Brogaard *et al*., 2012). Heatmaps showing +/−1kb regions around annotated nucleosome dyads. Average predictions are shown as metaprofiles on top of each heatmap. Plots from *left* to *right* display predictions of the following models: *in vitro* nucleosome ChromWave, *in vivo* nucleosome ChromWave, and nucleosome prediction and TF predictions from the joint TF-nucleosome ChromWave model. Each track was normalised by subtracting the genomic average before plotting. **D.** Motif detectors associated with *in vivo* nucleosome occupancy. Each point in the scatter plot represents a motif that is PCA embedded using maximum activations of motif detectors per sequence as input. The shape of each point indicates if the motif was annotated by TOMTOM (triangle) or de novo discovered (point). The size of each point was scaled with the mean activation of the motif detector. The colour scale corresponds to the mean Pearson correlation between activation and prediction per base-pair. Selected motifs were annotated with their associated sequence logo. **E.** *In vivo* and *in vitro* nucleosome occupancy prediction per base-pair centered on TF binding sites defined by ChIP-exo. Heatmaps showing +/−1kb regions around the ChIP-exo peaks. Each heatmap panel is based on a different TF peak set from top to bottom: Abf1, Cbf1, Rap1 and Reb1. Heatmaps from *left* to *right* display predictions of the following models: *in vitro* nucleosome ChromWave, *in vivo* nucleosome ChromWave, and nucleosome prediction and TF predictions from the joint TF-nucleosome ChromWave model. Average predictions (logfc-RPM for the nucleosome-only model, FPM for TF-nucleosome model) are shown as metaprofiles on top of each heatmap. Each track was normalised by subtracting the genomic average before plotting.

To test the accuracy of the models in predicting nucleosome positions, we defined ‘enriched’ and ‘depleted’ regions following (Kaplan *et al*., 2009) ChromWave’s *in vivo* and *in vitro* models were able to distinguish enriched and depleted regions defined from the *in vivo* MNase-seq data better than Kaplan HMM (Kaplan *et al*., 2009)(measured as AUC, the values were 0.99, 0.94 and 0.93, respectively; Figure S2C). Both predicted signal amplitudes and spacing between enriched regions matched those of the observed *in vivo* MNase-seq data (Figure S2D).

Finally, we aligned the ChromWave predictions to the nucleosome dyad positions defined by chemical cleavage (Brogaard *et al*., 2012) or ATAC-Seq (Schep *et al*., 2015). Both models learned the strongly positioned nucleosome at the dyad: the *in vivo* model produced a phasing pattern on either side of the central nucleosome that is similar to the TF+nucleosome ChromWave model (Figures 6C and S3A). However, the *in vitro* model failed to capture the phasing pattern. This is in line with the original MNase-seq data (Figure S3B) and previous experimental observations that phasing is lost if certain factors such as remodelers RSC and SWI/SNF and general regulatory factors Abf1 and Reb1 are absent (Gkikopoulos *et al*., 2011; Zhang *et al*., 2011; Krietenstein *et al*., 2016). This is also reflected in the accuracy of the NFR predictions around TSSs: this characteristic depletion in average nucleosome occupancy *in vivo* is correctly predicted by the *in vivo* ChromWave model, together with the phasing of nucleosome both up- and downstream of the NFR in contrast to both the HMM and the *in vitro* ChromWave model (Figure S2B).

### 2.8 Determinants of *in vivo* nucleosome occupancy encoded in the DNA sequence

The *in vivo* ChromWave model uses the pre-trained filters of the *in vitro* model as fixed convolutional filters; thus the trainable parameters have been used to learn subtle differences between the *in vivo* and *in vitro* nucleosome maps. To understand what additional information the *in vivo* model has learned, we inspected the convolutional filters that are unique to this model compared with the *in vitro* one.

We ranked the 256 *in vivo*-specific motif filters according to their activation, computed using their maximum output values (maximal activation) for each 2Kb DNA-sequence input (see *‘Convolutional Layers’* in Figure 1 where the output of the filters when applied to a one-hot encoded DNA-sequence is annotated as a blue square before being concatenated). The maximal activation provides a position-independent measure of the occurrence of the learned motifs (or highly similar motifs) for each DNA input sequence. To interpret the impact of motifs on ChromWave’s nucleosome predictions, we calculated correlations between the vectors of filter activation values for an input sequence and the predicted nucleosome occupancy. A positive correlation indicates that the presence of a sequence pattern matching the motif is favourable to nucleosome occupancy, whereas a negative correlation indicates that it is unfavourable. We used the mean of these correlations across all input sequences to interpret the effect of each filter across the whole genome.

In Figure 6D, we show a PCA plot of the maximal activations of the *in vivo*-specific filters across all DNA-sequence windows, with each point coloured according to their correlations. Many of the learned motifs occur only in a fraction of the input windows, whereas nucleosomes cover a large proportion of the genome. ChromWave’s architecture enables nucleosome occupancy prediction to be influenced by combinations of many different motifs, each a low individual impact by combining the output of individual filters many times. This is reflected by the moderate range of correlations for many individual filters in Figure 6D, rather than a dominating effect of only a few filters on the output.

We can see that motifs with similar GC-content tend to co-occur in the same input sequences with similar maximal activations. There are two large clusters: AT-rich motifs which are negatively correlated with nucleosome occupancy, and GC-rich motifs that positively correlate with occupancy. Among the AT-rich group are known motifs for sequence-specific chromatin remodelers such as Stb3, Sfp1, Dig1, Rlm1, and several TATA-box motifs. Poly-(d:A,d:T) stretches have been reported to deter nucleosome formation due to the rigidity of the DNA and thus are predictive of nucleosome binding (Segal and Widom, 2009). GC-rich filters include motifs for TFs whose DNA-binding correlate with nucleosome occupancy (for example Hal9, Cbf1, and YLL054C; (Charoensawan *et al*., 2012) and other motifs predominantly found in the 5’ NFRs, echoing many of the patterns learned by the TF+nucleosome ChromWave model (for example Gal80, Cbf1, and Pho2) (Figure 6D).

To compare the learned motifs across the different ChromWave models more systematically, we computed similarity scores and clustered the convolutional filters (excluding the pre-trained filters of each model) as previously described by Vierstra (Vierstra *et al*., 2020); Supplementary Table 2). 63 of the *in vivo-*specific 256 filters comprise similar motifs to those in the TF+nucleosome ChromWave model. Of the other 183 *in vivo-*specific filters, only 2 clustered with similar motifs found in the *in vitro* filters; both reflected RSC motifs. The other *in vivo*-specific filters included an additional RSC motif as well as sub-motifs of PHO4/RTG and MCM1/PDR1 (Figure S4).

Given the disparities in learned motifs, we hypothesized that the *in vitro* ChromWave model would not be able to correctly model nucleosome eviction at binding sites of general regulatory factors, whereas the *in vivo* model would be able to model this competition correctly. Indeed, when we aligned the predictions again around known binding sites of the Abf1, Rap1, Reb1 and Cbf1(Rossi, Lai and Pugh, 2018) we observed a stark difference between the predictions of the two ChromWave models: the *in vitro* model predicts a nucleosome directly on the binding sites of the TFs, while the *in vivo* model was able to learn nucleosome depletion on the binding sites, albeit not as well as the ChromWave model trained jointly on TF and nucleosome data (Figure 6E).

In summary, although ChromWave when trained on *in vivo* nucleosome occupancy maps can recover *in vivo* nucleosome positions to high accuracy, the missing information on TF binding hinders modelling the competition of nucleosomes with other sequence-driven binding events.

### 2.9 Predicting nucleosome occupancy in the human genome with ChromWave

Nucleosome occupancy models trained on yeast data have been reported to produce predictions that correlate well with nucleosome maps from other organisms (Gupta *et al*., 2008; Awazu, 2017). However, there are marked differences between the positioning of human and yeast nucleosomes given the same underlying DNA-sequence. When we applied the *in vitro* and *in vivo* ChromWave models to 23,156 human promoter regions, they produced terrible agreement with MNase-seq datasets from several lymphoblastoid cell lines(Gaffney *et al*., 2012; Zhao *et al*., 2018), with average Pearson correlation coefficients of 0.002 and −0.0003, respectively.

We, therefore, trained a new ChromWave model to predict *in vivo* nucleosome occupancies around human promoter sequences (+/−1KB around TSSs; Methods). We used transfer learning across organisms: the first convolutional layers of a yeast *in vivo* nucleosome occupancy ChromWave model was used as a set of pre-trained convolutional layers. The final model had only 32 additional trained convolutional layers and achieved excellent accuracy at class level (Figure S5A).

When mapping classes back to the original continuous values of the binned reads, ChromWave’s ability to achieve accurate predictions across all promoters with similar performance across the training, test, and validation sets is evident (Figures 7A and S6A). To measure the model’s ability to predict nucleosome positions correctly, we defined nucleosome enriched and depleted regions from predicted and observed occupancy profiles (Methods). ChromWave achieved an AUC of 0.63 when predicting enriched vs depleted regions. The predicted signal amplitudes and spacing between enriched regions matched those of the observed *in vivo* MNase-seq data (Figure S5B).

**Figure 7.**
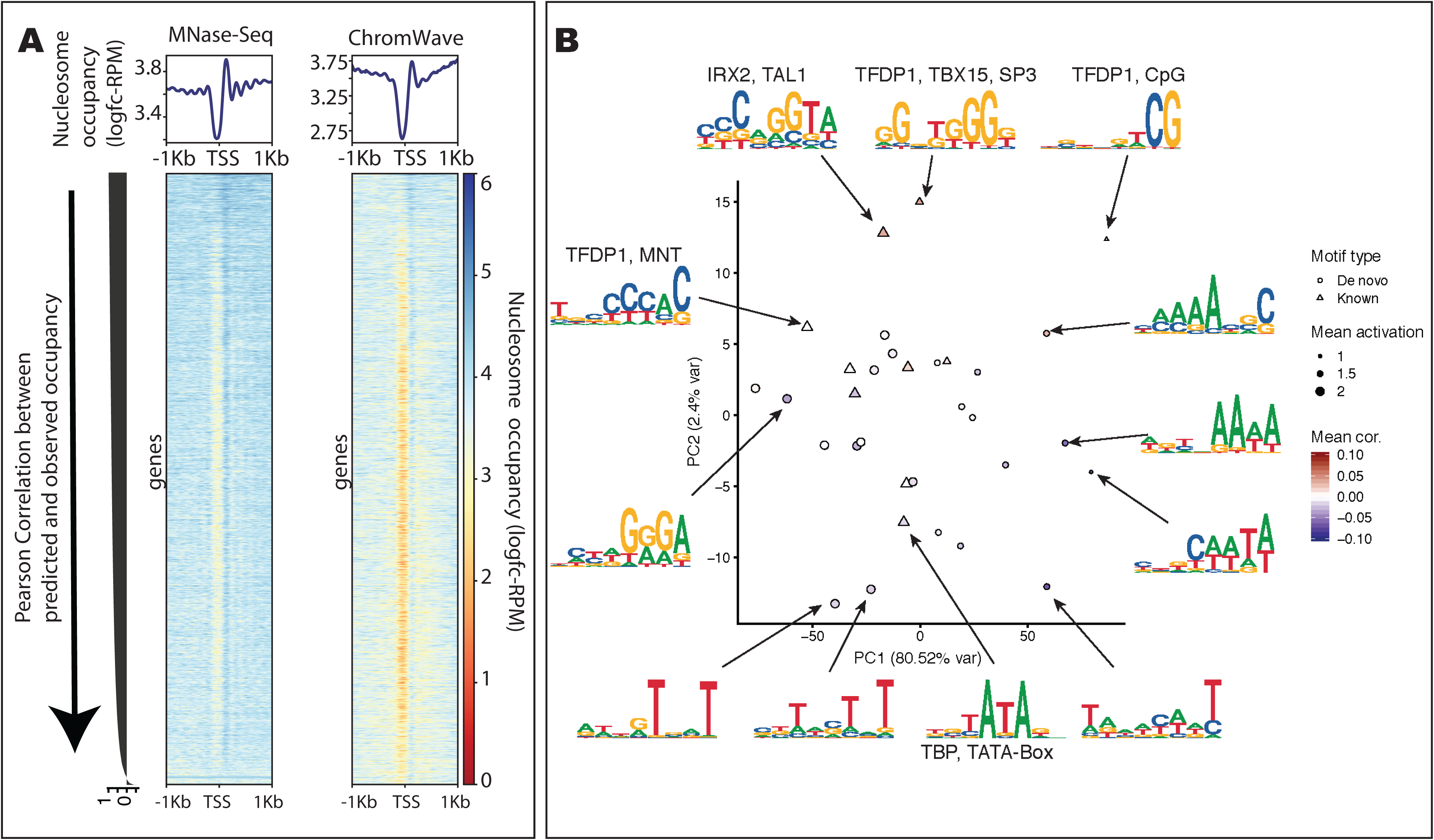
Predicting nucleosome occupancy in human promoters with ChromWave. **A.** Comparison of MNase-seq and predicted nucleosome occupancy in +/− 1kb around 23,156 annotated human TSSs. Occupancy is measured and predicted as log2-fold change of number of reads at each position over genome average, in Reads Per Million (logfc-RPM). *Top:* Metaprofiles of average nucleosome occupancy measured by MNase-seq (left) and predicted by ChromWave (right) are centered around TSSs. Heatmaps are showing the data and predictions of each TSS in the rows. TSS are sorted by the Pearson Correlation coefficients between observed and predicted profiles from top to bottom (as indicated by the arrow) and these are shown on the left as barplot. **B.** Motif detectors associated with human promoter nucleosome occupancy. Each point in the scatter plot represents a motif that is PCA embedded using maximum activations of motif detectors per sequence as input. The shape of each point indicates if the motif was annotated by TOMTOM (triangle) or de novo discovered (point). The size of each point was scaled with the mean activation of the motif detector. The colour scale corresponds to the Pearson correlation between activation and prediction per base-pair. Selected motifs were annotated with their associated sequence logo.

### 2.10 ChromWave learns known and unknown motifs associated with nucleosome binding and DNA-methylation in human promoter regions

Again we visualised the sequence motifs learned by the first convolutional layer and compared them to known motifs (JASPAR CORE vertebrates 2016, JASPAR POLII and HOCOMOCO v.10). We then repeated the PCA and correlation analysis described above (Section 2.8) to elucidate the co-occurrence of motifs and their impact on nucleosome occupancy predictions. As for the yeast *in vivo* model, ChromWave learned AT-rich motifs, including the TATA-box binding motif, that anti-correlate with nucleosome occupancies in human promoters, and GC-rich motifs that are positively correlated, including a CpG pattern (CG) which is associated with DNA methylation in humans (Schübeler, 2015), and the motifs of TFDP1, SP3, and MNT (Figure 7B), three proteins that are excluded upon methylation of open chromatin (Bartke *et al*., 2010).

We hypothesized that a model trained only on promoters might lack the information necessary to predict nucleosome occupancies in other types of genomic regions. For instance, CTCF-binding sites in intergenic regions are known to be depleted of nucleosomes (Kim *et al*., 2007). We asked ChromWave to make predictions around experimentally identified CTCF sites (Davis *et al*., 2018; Zhang *et al*., 2020), and in stark contrast to the MNase-seq data, the model placed nucleosomes directly over the CTCF sites irrespective of the binding strength of CTCF (Figure S6B).

### 2.11 Genetic components of promoter architecture in the human genome

To assess the impact of sequence changes at human promoters, we used ChromWave to prioritise single nucleotide polymorphisms (SNPs) by their predicted impact on nucleosome positioning.

First, we predicted the effects of sequence changes at *DNase I sensitivity quantitative trait loci* (dsQTLs; 1366 of 6057 fell within +/−1kb around TSS; (Degner *et al*., 2012). These are genetic variants that modify chromatin accessibility as measured by DNase I sequencing. Although many dsQTLs did not affect predictions, we identified 30 promoters where ChromWave’s chromatin accessibility predictions based on MNase-seq data were highly affected by dsQTLs (Figure 8A and Supplementary Table 3). We used the annotation of DNase I hypersensitive sites (DHS) from the same study (Degner *et al*., 2012) to annotate SNPs inside and outside of DHSs. Of the 30 dsQTLs that we identified to change chromatin accessibility, 7 were inside an annotated DHS, whereas the rest acted distally.

**Figure 8.**
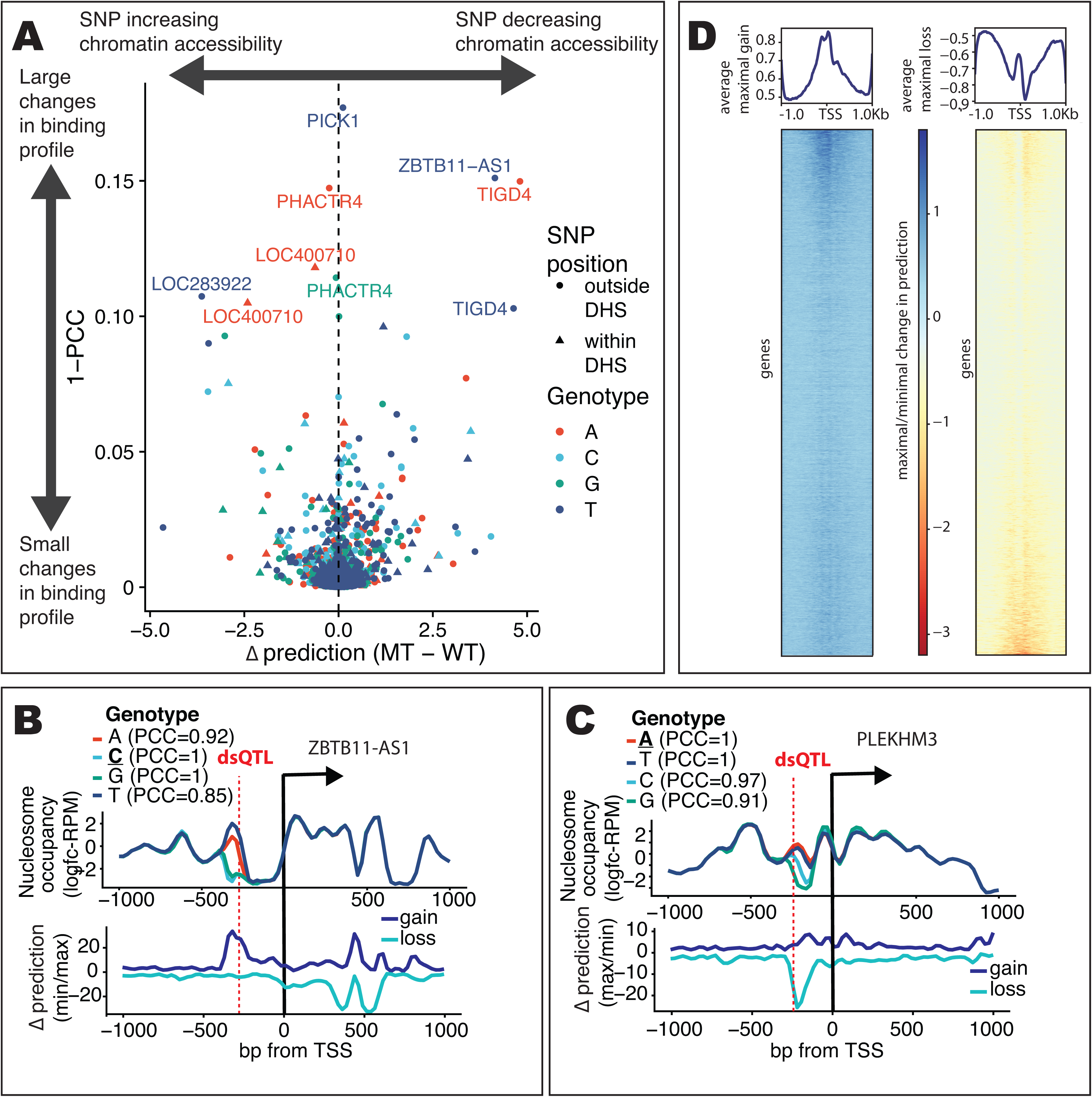
ChromWave learns known and unknown motifs associated with nucleosome binding and DNA-methylation in human promoter regions. **A.** Volcano plot to identify dsQTLs that change nucleosome occupancy in promoter regions. Each point represents a dsQTL variant, some annotated with a promoter name. Along the x-axis, we plot the difference (Δ prediction (MT-WT)) between the predicted nucleosome occupancy given dsQTL variant and the wild-type input sequence (e.g. the ratio between the predicted raw read counts between the signle nucleotide change and wild type). Along the y-axis, we show (1-the Pearson correlation coefficient) between the predicted nucleosome occupancy given the dsQTL variant and the wild-type input sequence. The shape of each point indicates if dsQTL falls within or outside of the previously associated DNase I hypersensitive site (DHS). The colours represent the genotype of the variant as indicated. **B.** Example of maximal disruptive dsQTLs on nucleosome occupancy predictions. *Top Panel:* Nucleosome occupancy for different genotypes at the dsQTL (indicated by the dotted red line) compared with observed data. The reference genotype “C” is indicated in bold. *Bottom Panel:* Line plots show the maximal increase (‘gain’) and maximal decrease (‘loss’) in the predictions at each position given all possible single base-pair changes along the sequence. **C.** Example of maximal disruptive dsQTLs on nucleosome occupancy predictions. As B.The reference genotype “A” is indicated in bold. **D.** Hotspots in human promoters susceptible to changes in chromatin accessibility due to single nucleotide polymorphisms. Maximal increase (‘gain’) and maximal decrease (‘loss’) in the predictions at each position given all possible single base-pair changes along the sequence in +/− 1kb around 23,156 annotated human TSSs. *Top:* Meta-profiles of maximal increase (‘gain’) (*left*) and maximal decrease (‘loss’) (*right*) in ChromWave’s predictions centered around the TSSs. *Bottom:* Heatmaps are showing the maximal increase and decrease for each TSS in the rows.

We focused on dsQTLs that changed ChromWave’s prediction locally at the dsQTL itself (Δ prediction SNP-WT > 0 or Δ prediction SNP-WT < 0), as well as a dramatic effect on the overall nucleosome occupancy profile measured by the Pearson correlation coefficient (PCC) between the predicted WT and SNP-perturbed profiles (1-PCC>0.05). Another way of interpreting the scale of the x-axis is as the ratio between the predicted raw read counts between the mutant and wild-type since both profiles are predicting the log2-fold change between reads per million and the total average (and log(x/N)-log(y/N)=log(x/y)). We identified 17 dsQTLs leading to gains in nucleosomes thus decreasing chromatin accessibility (1-PCC > 0.05 and Δ prediction SNP-WT > 0) and 13 dsQTLs causing losses (1-PCC > 0.05 and Δ prediction SNP-WT < 0).

The promoter of the ZBTB11-AS1 gene displays one of the largest changes in both the local prediction and the overall binding profile (1-PCC > 0.15, Δ prediction [C>T] - WT > 4). The predictions of the promoter sequence with different bases substituted at the dsQTL are shown in Figure 8B. The reference genotype (C) and a C>G change at the dsQTL yield the lowest predicted local nucleosome occupancy at the dsQTL, whereas C>A and C>T changes cause dramatic increases in predicted occupancies, with a strongly positioned nucleosomes directly on the dsQTL (Figure 8B, top panel). The PLEKHM3 promoter is an example of a loss in nucleosome occupancy following a single nucleotide change (Figure 8C): alteration from the reference genotype (A) to C or G at the dsQTL removes a weakly positioned nucleosome (1-PCC >0.09, Δ prediction [A>G] - WT < −3). An A>T change does not alter the profile (Figure 8C, top panel).

### 2.12 Changes in nucleosome-binding in human promoters induced by simple sequence changes are restricted to ‘hot spots’ which cover previously identified dsQTLs

To assess the impact of sequence changes in the whole promoter region on nucleosome occupancy, we computed the maximal gain and loss in prediction at each position given all possible base changes along the sequence for all promoter regions in the dataset.

The bottom panels in Figures 8B and 8C show these maximal gains and losses for the two example promoters of ZBTB11-AS1 and PLEKHM3. In both promoter regions, not all nucleosome position predictions are dynamic under sequence changes. Instead, the predicted decrease and increase in prediction is restricted to small regions in the promoters (some of which coincide with the position of the dsQTL), while most nucleosome occupancy predictions are robust to individual base changes.

These observations suggest that most nucleosome positions in human promoter regions are robust to sequence changes while some can be altered considerably through simple base changes. Figure 8D shows individual and average profiles of maximal gains and losses of nucleosome occupancy prediction given all possible base-changes in all promoter regions in the dataset. The nucleosome occupancy of around two-thirds of human promoters is affected by single nucleotide changes in 1000bp around the TSSs in either gain or loss (Figure 8D). These changes of occupancy are indeed not randomly distributed in the promoters. Instead, given the right single nucleotide changes (compared with wild-type), more nucleosomes bind close to and directly on the TSS (Figure 8D left). However, given the right single nucleotide changes (compared to wild-type) nucleosomes are lost just up- and downstream of the TSS, while nucleosome occupancy and depletion directly on the TSS and in the NFR just upstream of the TSS are less affected (Figure 8D right).

## 3 Discussion

We presented a deep learning framework called ChromWave to model DNA-binding profiles of nucleosomes and TFs simultaneously at nucleotide resolution to high precision and deployed it genome-wide in yeast across different data sets and experimental methods as well as in human promoter regions. ChromWave learned intricate sequence grammar of DNA accessibility and competition between DNA-binding proteins. ChromWave surpasses with its precision, scope, and flexibility current state-of-the-art methods to predict TF and nucleosome binding from DNA-sequence alone. We demonstrated and visualised how ChromWave identifies and uses specific sequence features that encode information for DNA-binding and the competition between nucleosomes and TFs.

We showed that ChromWave recapitulates computationally results from a previous study that was based on lab experiments (Floer *et al*., 2010) to determine TF and nucleosome binding in the well-studied GAL1-GAL10 locus. Taken together, our results match the experimentally derived interpretation in (Floer *et al*., 2010): a small (partially unwound) nucleosome is positioned at the locus UASg by the RSC complex thereby facilitating Gal4 binding to its side. Upon deletion of the RSC binding site, nucleosomes encroach over the UAS and compete with Gal4 for binding. These results illustrate how ChromWave can be used to generate and test hypotheses *in silico* before testing these in the lab.

In the human genome, ChromWave lets us annotate dynamic hotspots where nucleosome occupancy is highly affected by simple base changes in the DNA sequence compared with the rest of the promoter region. By comparing models that were trained on different types of data, we have qualified sequence patterns that allow ChromWave to correct *in vitro* nucleosome positions to *in vivo.* In addition, these experiments have shown that lacking TF-binding information severely impedes the capability of the model to predict nucleosome positions to high precision and the models are unable to capture the competition between different DNA-binding factors.

Previous work to model TF- and nucleosome-binding at nucleotide-resolution has been constrained to the usage of HMMs in the yeast genome (Wasson and Hartemink, 2009; Ozonov and van Nimwegen, 2013; Zhong, Wasson and Hartemink, 2014). An advantage of these models over ChromWave is that they can resolve which TF is predicted to be bound at a certain position based on the assumption that the TF has a well-annotated motif. However, due to the computational complexity of determining concentrations for all used TFs in the model, these studies had to constrain their models to only a small set of TFs (158 TFs were used in (Ozonov and van Nimwegen, 2013), 89 in (Wasson and Hartemink, 2009), and only 42 in (Zhong, Wasson and Hartemink, 2014)). Nevertheless, our work was inspired by these early studies; in fact, we used the same training data and similar pre-processing methods as in (Zhong, Wasson and Hartemink, 2014) albeit the former study was constrained to promoter regions. At the expense of the ability to distinguish different TF binding events, ChromWave can predict nucleosome- and TF-binding genome-wide, and can resolve binding events not directly explainable by an annotated motif.

Our work also builds on the success of recent deep learning methods that model DNA-binding and gene expression, e.g. (Alipanahi *et al*., 2015; Kelley, Snoek and Rinn, 2015; Quang and Xie, 2016; Kelley *et al*., 2018). Most of these models perform binary classification tasks and predict whether a relatively short DNA-sequence (30bp-1000bp) will be bound by a TF or RNA-binding protein (Alipanahi *et al*., 2015), a nucleosome (Quang and Xie, 2016), or surrounds a DNase I hypersensitive site in different cell types (Kelley, Snoek and Rinn, 2015). None of these models predict two competing binding events at the same time, nor do they make a prediction for each of the input positions and instead summarise of the whole sequence. In contrast, (Kelley *et al*., 2018) developed the first deep learning model, called Basenji, that can model distal regulatory interactions to predict quantitative (as opposed to binary) genomic profiles such as read counts of DNase I seq, histone modification ChIP-seq, and expression data. ChromWave takes a similar approach by modeling read counts of MNase-seq data. Basenji can take 131kb regions as input which is much larger than the 2000bp-5000bp we chose for ChromWave, however this scale comes at the cost of resolution: while Basenji (and its recent extension the Enformer (Avsec, Agarwal, *et al*., 2021)) predicts one value for every 128bp bins (which already was a refinement over the 200bp bins of ChromHMM (Ernst and Kellis, 2017)), ChromWave makes a prediction for each individual nucleotide. This is an important improvement as most of our results on the intricate competition of TFs and nucleosomes display changes on just a few base pairs with most TF binding sites predicted to be shorter than 15bp (Stewart, Hannenhalli and Plotkin, 2012). This resolution has only been achieved by a recently published model (BPNet) which shares many similarities in its architecture with that of ChromWave (Avsec, Weilert, *et al*., 2021). While BPNet predicts several TF binding profiles simultaneously and the study focused on TF binding motifs, nucleosome occupancies and their competition was largely ignored.

The main limitation of any model relying on training data restricted to certain regions in the genome is illustrated by our human promoter ChromWave model predicting nucleosomes at CTCF-sites: as we only trained in promoter regions, its genome-wide predictions cannot be trusted as it was not exposed to binding factors outside of promoter regions. To remedy this bias, ChromWave could be further trained on more DNA sequences that span the (mappable) genome. Another approach may be using an ensemble of several ‘expert’ ChromWaves models that have learned specific sequence context (e.g ‘promoter regions’, ‘insulator sites,..). Either of these approaches can be realised with the ChromWave framework, however, this observation highlights the importance of understanding the data and training details whenever models such as ChromWave (and their predictions) are being used.

Although we presented ChromWave as a model of different chromatin accessibility profiles, ChromWave is a flexible framework to model genome-wide chromatin-modification. as well as transferable to mammalian genomes. As such, ChromWave is an unprecedented framework to study how genetic variability impacts DNA-binding. Although not demonstrated in this study, profile-like signals such as histone marks and DNA-methylation could form the prediction target to ask ChromWave to predict these signals simultaneously *in silico* at nucleotide resolution. Other applications of ChromWave may be modelling time-dependent chromatin profile changes, where we expect the environmental changes to have an effect on DNA-binding. As a proof of concept, we have successfully used ChromWave to model nucleosome profiles of young and old yeast (data not shown, but code and data are available on github https://github.com/luslab/ChromWave). Other applications of ChromWave may be the modelling of binding profiles during differentiation, or hormone changes.

In summary, ChromWave allows the prediction of TF and nucleosome binding simultaneously genome-wide at nucleotide resolution with high precision. The sequence-based approach allows the annotation of every mutation in the genome that influences chromatin accessibility through TF or nucleosome binding. ChromWave has the potential to provide important insights into the effect of genetic variation on transcriptional changes in complex diseases, at a certain time-point, but also during changes over time. As more and more non-coding variants are discovered in the human genome, these insights can provide us with an important understanding of how SNPs affect biological mechanisms and phenotypes.

## Acknowledgements

This research was funded in whole, or in part, by the Wellcome Trust (FC010110; 215593/Z/19/Z). For the purpose of Open Access, the author has applied a CC BY public copyright license to any Author Accepted Manuscript version arising from this submission. This work was supported by the Francis Crick Institute which receives its core funding from Cancer Research UK (FC010110), the UK Medical Research Council (FC010110), and the Wellcome Trust (FC010110). NML is a Winton Group Leader in recognition of the Winton Charitable Foundation’s support towards the establishment of the Francis Crick Institute. NML is additionally funded by a Wellcome Trust Joint Investigator Award (215593/Z/19/Z) and core funding from the Okinawa Institute of Science & Technology Graduate University.

We thank Joe Brock (The Francis Crick Institute) for helping with illustrations, and Drs. Dean Plumbley (BenevolentAI), Joe Ledsam (Google DeepMind, Google Japan), Bernadino Romera-Paredes (Google DeepMind), Erik Pfeiffenberger (The Francis Crick Institute, BMW Group), and Prof. David Jones (The Francis Crick Institute, UCL) for many interesting discussions and their valuable advice and helpful comments regarding deep learning, neural networks, and visualisation techniques.

## Author Contributions

Conceptualization, S.A.C. and N.M.L.; Methodology, S.A.C. and S.S.; Software, S.A.C, S.S, J.S and W.X.; Validation, S.A.C and S.S; Formal Analysis, S.A.C., S.S. and N.M.L.; Investigation, S.A.C. and S.S.; Resources, W.X. and N.M.L; Writing – Original Draft, S.A.C. and N.M.L.; Writing – Review & Editing, S.A.C. and N.M.L; Visualization, S.A.C. and S.S.; Supervision, S.A.C. and N.M.L.

## Declaration of Interests

The authors declare no competing interests.

## Figure Legends

**Supplementary Figure 1.**
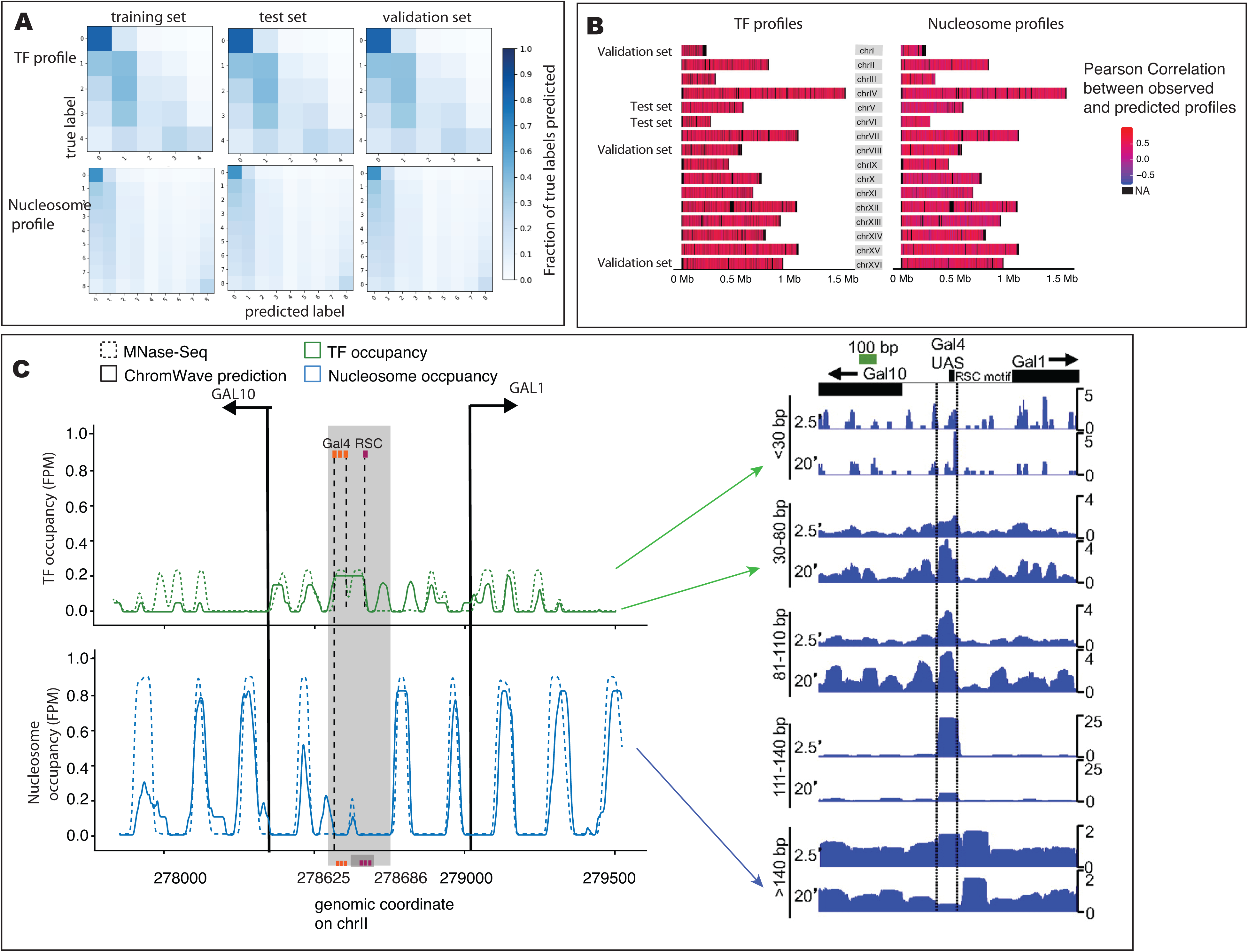
**A.** Normalised confusion matrices comparing the true classes and the predictions for TF and nucleosome profiles computed on the training, test and validation sets as indicated. The matrices were normalised by the number of elements in each class, e.g. the colour of tile (i,j) represents the fraction of true labels i that were predicted to be class j and each row sums to 1. **B.** Genome-wide comparison ChromWave’s predictions of TF and nucleosome occupancy and observed data. Shown are Pearson Correlation coefficients between predicted and observed TF (left) and nucleosome (right) occupancies (in Fragments Per Million, FPM). The chromosomes forming the validation and test sets are indicated (all other chromosomes formed the training set). Correlations were computed in sliding windows of size 300bp and averaged across 300bp bins for visualisation. **C.** *Left:* MNase-seq profiles and ChromWave’s predictions (in Fragments Per Million, FPM) of TF (green) and nucleosome (blue) occupancy in the intergenic region on chromosome II 2 between the GAL10 and GAL1 genes on chromosome II. Annotations of Gal4 and RSC motifs are indicated by stippled lines. The light shaded region indicates a yeast regulatory locus GAL4 UAS. Inside, a partially unwrapped nucleosome bound by RSC is shown as a broad peak in the TF profile as the nucleosome is represented by shorter reads. *Right:* Original Figure 3A from (Henikoff *et al*., 2011) for comparison. Shown are mapped MNase-Seq reads at different digestion times (2.5 and 20 min). Arrows from the left figure to the right figure indicate the equivalent read tracks (at 20min digestion time). The partially unwrapped nucleosome apparent in the TF profile on the left is apparent in the larger fragments (>140bp) at short digestion time (2.5 min).

**Supplementary Figure 2.**
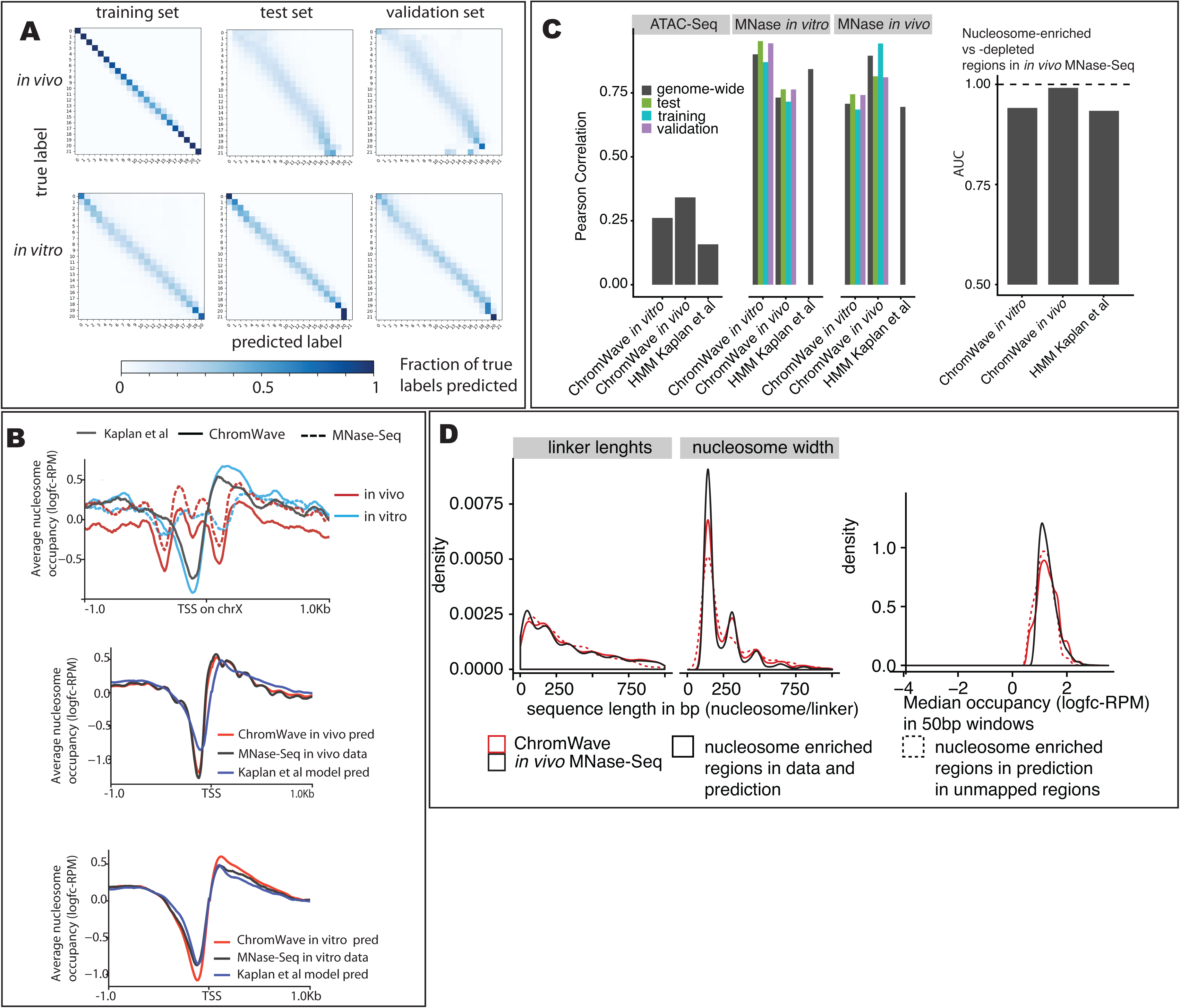
**A.** Normalised confusion matrices comparing the true classes and the predictions for *in vitro* and *in vivo* nucleosome profiles computed on the training, test and validation sets as indicated. The matrices were normalised by the number of elements in each class, e.g. the colour of tile (i,j) represents the fraction of true labels i that were predicted to be class j and each row sums to 1. The highest class (21) was very rare in the in vitro data (observed at ~400bp genome-wide) and did not occur in the training data. **B.** Comparison of nucleosome occupancy across models and across in vivo and in vitro datasets in yeast promoter. Metaprofiles +/− 1bp around all TSS on chromosome 10 of average predicted and observed nucleosome occupancy (in logfc-RPM)(upper panel). Metaprofiles +/− 1bp around all TSS of average predicted and observed nucleosome occupancy (in logfc-RPM) *in vitro* (middle panel) and *in vivo* (lower panel). Each track was normalised by subtracting the genomic average before plotting. **C.** Genome-wide comparison of predicted and observed nucleosome occupancy across different datasets. *Left:* Shown are Pearson’s correlation coefficients of the predicted occupancy of the *in vitro* and *in vivo* nucleosome ChromWave models and the HMM from (Kaplan *et al*., 2009) with ATAC-Seq (Schep *et al*., 2015) and the *in vitro* and *in vivo* MNase-seq data both genome-wide and separately on training, test and validation sets, as indicated. *Right:* Area-under-the-ROC-Curve (AUC) measuring the ability of the models to distinguish nucleosome-enriched vs nucleosome-depleted regions in the different datasets. Enriched and depleted regions were defined as any window of 50bp with an average predicted occupancy exceeding 0.5 standard deviations above or below the genome-wide mean, respectively. Predictions were mapped back to the original (continuous) ranges before calculating enrichment. A score of 0.5 means the model is not performing better than chance, a value of 1 means perfect agreement with the data **D.** Left: Density plots of linker lengths and nucleosome widths for *in vitro* and *in vivo* nucleosomes (called as ‘enriched’ regions) from data and predictions in mapped and unmapped regions. Right: Density plots of signal amplitudes for *in vitro* and *in vivo* nucleosomes (called as ‘enriched’ regions) from data and predictions in mapped and unmapped regions. To plot signal amplitudes we applied a jitter of 0.4 to ChromWave’s predictions to smooth out the discrete scale for visualisation.

**Supplementary Figure 3.**
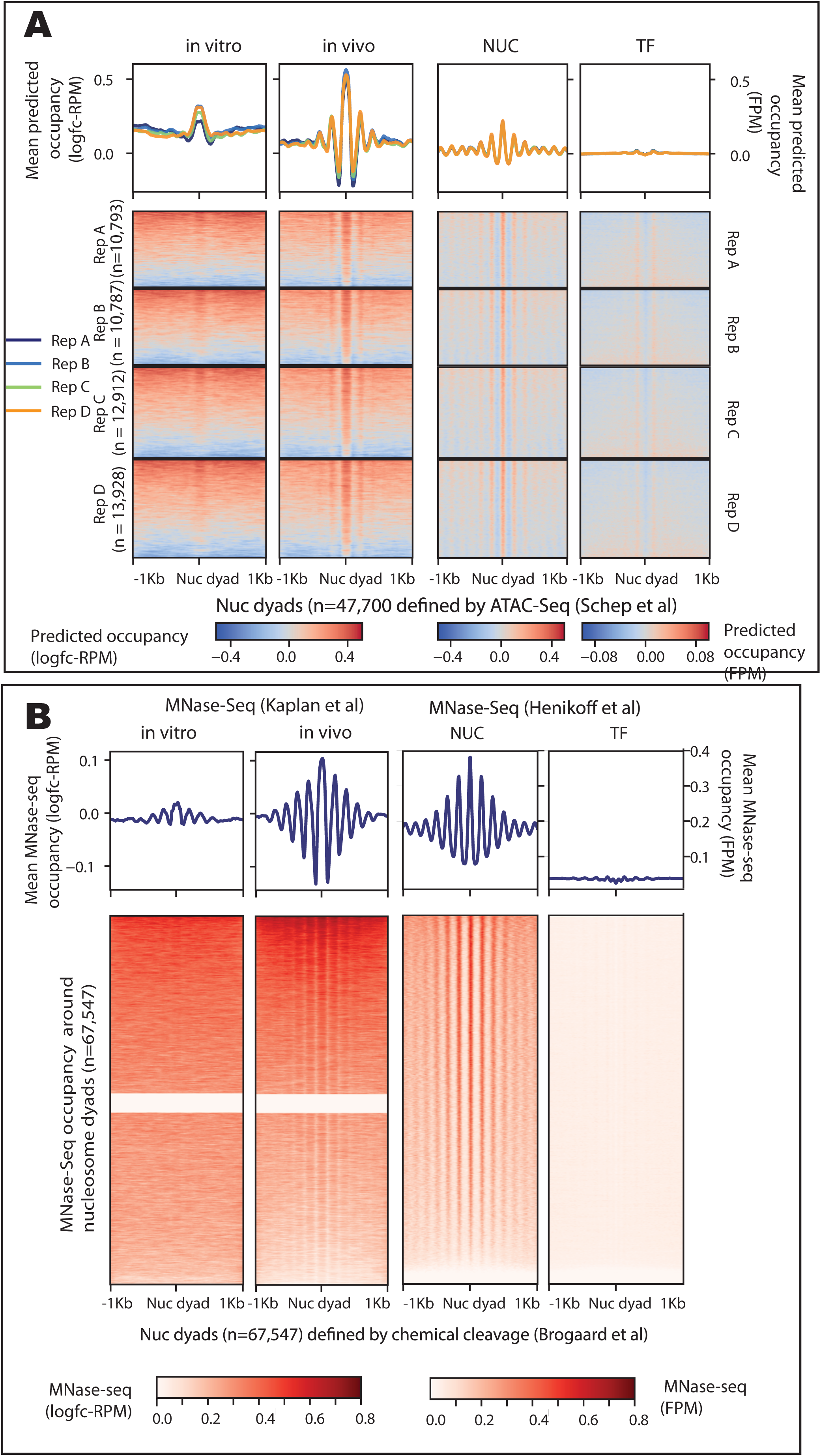
**A.** Model predictions per base-pair centered on nucleosome dyads called on ATAC-seq. Occupancy is measured and predicted as log2-fold change of number of reads at each position over genome average, in Reads Per Million (logfc-RPM). Heatmaps showing +/−1kb regions around ATAC-seq nucleosome dyads for four biological replicates (A, B, C, D). Average predictions (in logfc-RPM) are shown as metaprofiles on top of each heatmap. Plots are displaying nucleosome occupancy prediction using following models: *in vitro* nucleosome ChromWave (“*in vitro”)*, *in vivo* nucleosome ChromWave (“*in vivo”)*, and nucleosome prediction (“NUC”) and TF predictions (“TF”) from the joint TF-nucleosome ChromWave model. Each track was normalised by subtracting the genomic average before plotting. **B.** MNase-seq occupancy profiles centered on nucleosome dyads defined by chemical cleavage (Brogaard *et al*., 2012). Plots are displaying nucleosome and TF occupancy using following data: *in vitro* nucleosome (“*in vitro”)* and *in vivo* nucleosome (“*in vivo”)* from (Kaplan *et al*., 2009) (measured as log2-fold change of number of reads at each position over genome average, in Reads Per Million (logfc-RPM)), and nucleosome (“NUC”) and TF (“TF”) in Fragments Per Million (FPM) from (Henikoff *et al*., 2011). Heatmaps showing +/−1kb regions around nucleosome dyads. Dyads not covered by the nucleosome MNase-seq data from (Kaplan *et al*., 2009) show as no signal in the two left heatmaps. Average occupancies are shown as metaprofiles on top of each heatmap.

**Supplementary Figure 4.**
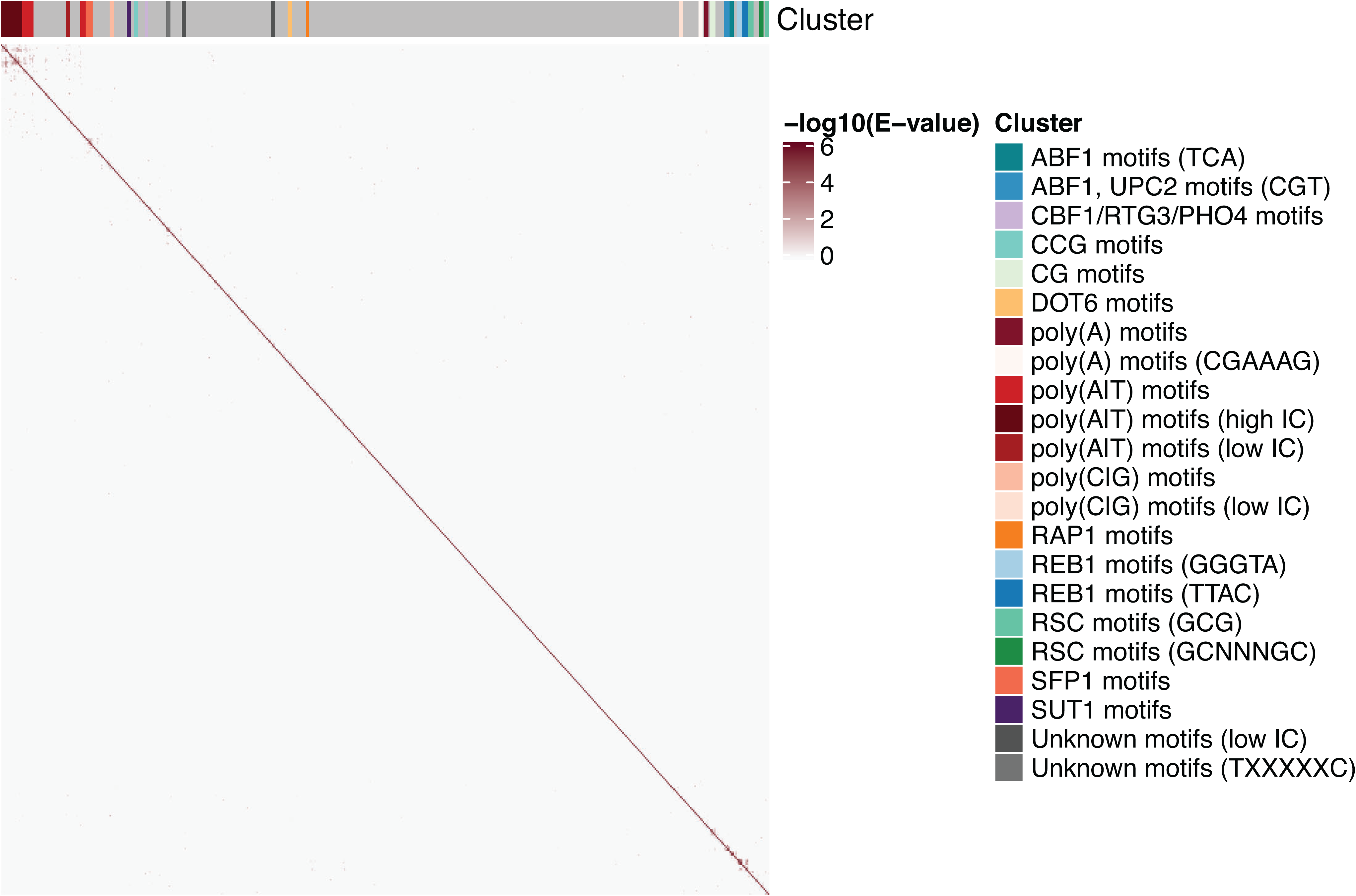
**A.** Motif clustering on the convolutional filter from all yeast models. Briefly, pairwise motif similarities were computed between the filters from all yeast models and used as input for hierarchical clustering visualised here as heatmap. Clusters were subsequently defined by cutting the dendrogram at height h=0.9 indicated in the barplot on top of the heatmap. The clusters were manually annotated using previous TOMTOM results as shown on the right.

**Supplementary Figure 5.**
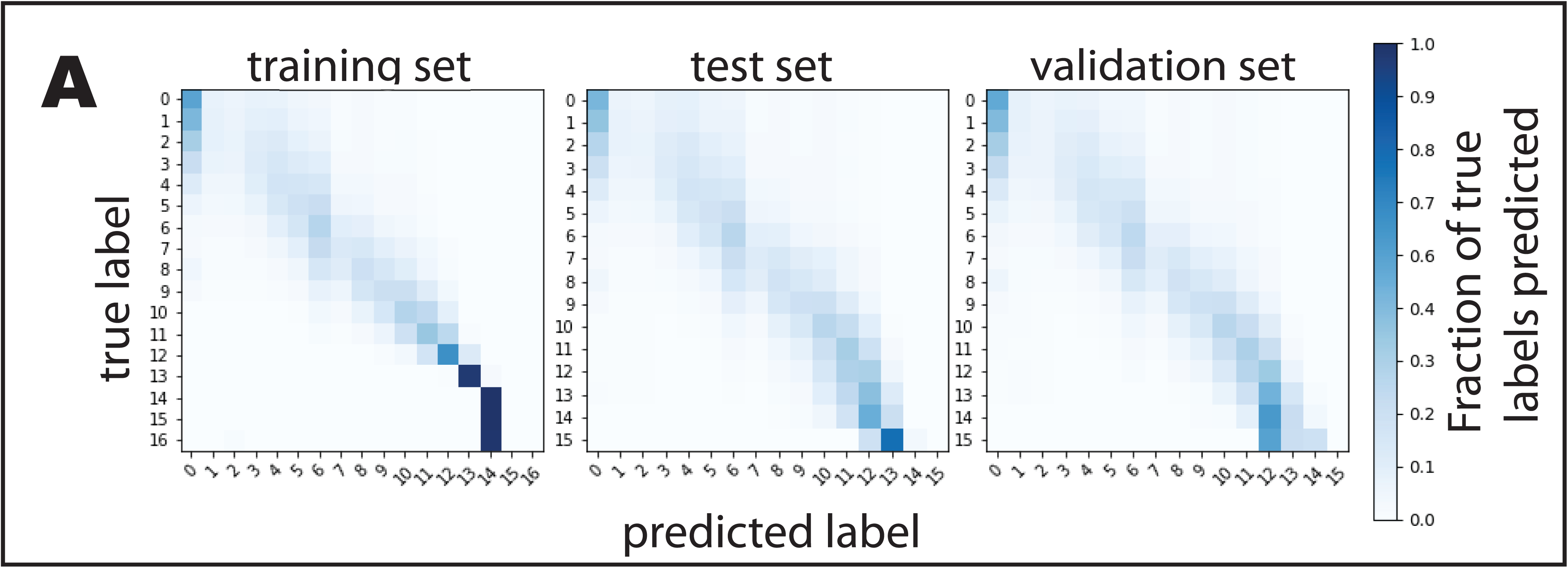
**A.** Normalised confusion matrices comparing the true classes and the predictions of nucleosome profiles around human TSSs computed on the training, test and validation sets as indicated. The matrices were normalised by the number of elements in each class, e.g. the colour of tile (i,j) represents the fraction of true labels i that were predicted to be class j and each row sums to 1.

**Supplementary Figure 6.**
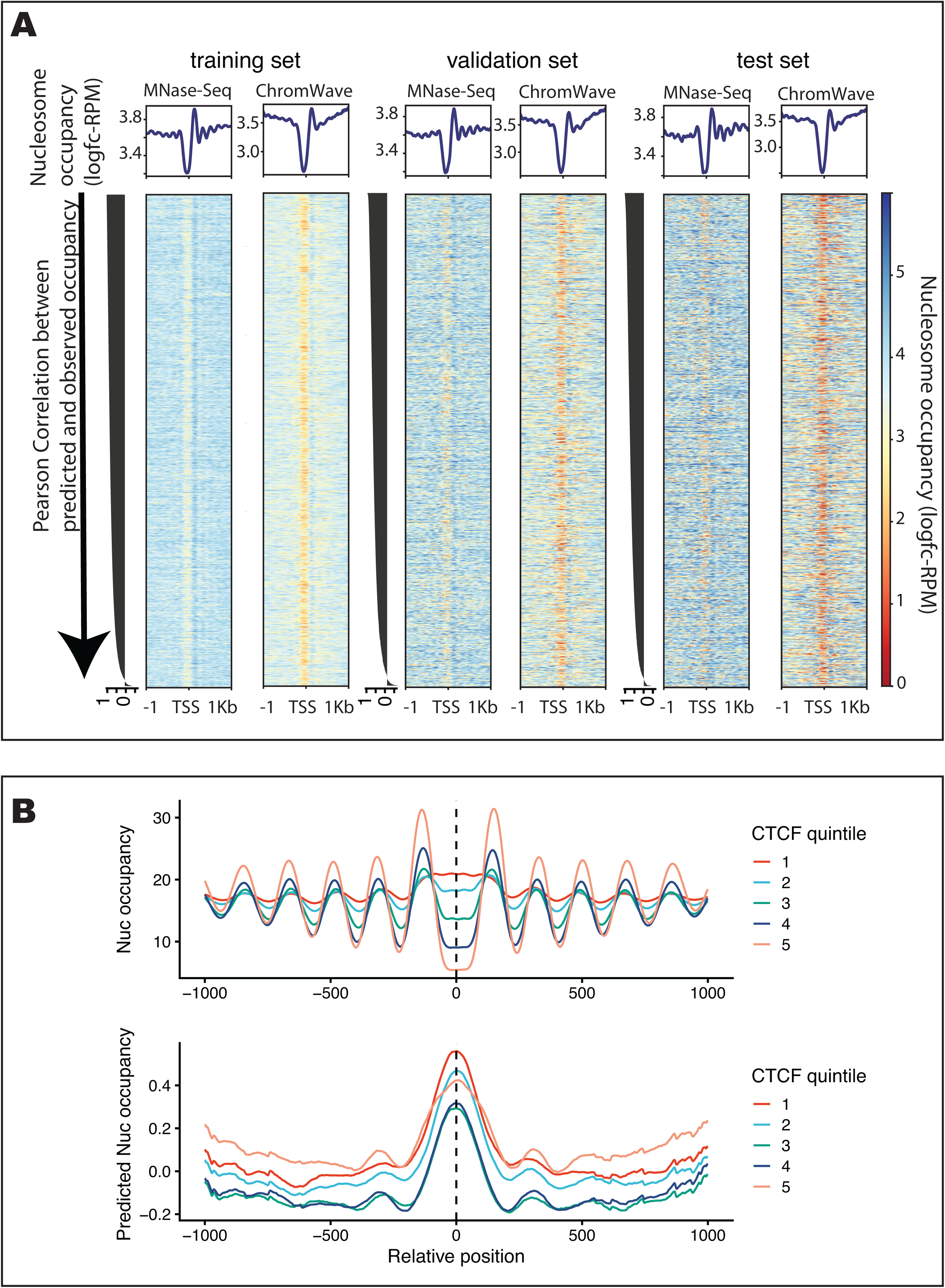
**A.** Comparison of MNase-seq and predicted nucleosome occupancy +/− 1kb around all human TSSs grouped by training, test and validation sets as indicated. Occupancy is measured and predicted as log2-fold change of number of reads at each position over genome average, in Reads Per Million (logfc-RPM). *Top:* Meta-profiles of average nucleosome occupancy measured by MNase-seq (left) and predicted by ChromWave (right) (in logfc-RPM) are centered around 23,156 annotated TSSs. *Bottom:* Heatmaps are showing the data and predictions of each TSSs in the rows. TSS are sorted by the Pearson Correlation between observed and predicted profiles from top to bottom as indicated by the arrow. Individual Pearson correlation coefficients are shown on the left of each heatmap as barplot. **B.** MNase-seq and predicted nucleosome occupancy around ChIP-seq CTCF binding sites. Metaprofiles are centered on CTCF peak summits, and these were split into quintiles according to the CTCF peak scores. *Top*: Metaprofile of the average nucleosome occupancy, as measured by MNase-seq (in logfc-RPM), for +/− 1kb around CTCF binding sites. *Bottom:* Metaprofile of ChromWave’s predicted average nucleosome occupancy (in logfc-RPM). Colors indicating CTCF quintiles, where quintile “five” corresponds to highest, and quintile “one” to the lowest ChIP-seq scores.

## Methods

### RESOURCE AVAILABILITY

#### Lead contact

Further information and requests for resources should be directed to and will be fulfilled by the Lead Contact, Dr Sera Aylin Cakiroglu.

#### Materials Availability

This study did not generate new unique reagents.

#### Data and Code Availability

The code, trained models, preprocessed input data and model predictions as BigWig files are publicly available on github (https://github.com/luslab/ChromWave). R-scripts implementing several visualisation methods can also be found on github (https://github.com/luslab/chromWaveR). The hyperparameter search space as well as the optimal parameters for all models can be found in Supplementary Table1. Motif clusters of the yeast models can be found in Supplementary Table 2. Human promoters with variable nucleosome occupancy at different base-changes in dsQTLs are listed in Supplementary Table 3.

### METHOD DETAILS

#### Data Preparation

##### Yeast data sets

We obtained the reference genome sequences of sacCer1 and sacCer3 (Engel *et al*., 2014). We used the *biomaRt* package (Durinck *et al*., 2005, 2009) to retrieve gene annotation information from Ensembl release 91 (Zerbino *et al*., 2018) using BioMart web services (Kasprzyk *et al*., 2004; Smedley *et al*., 2015). We downloaded the nucleosome occupancy maps from (Kaplan *et al*., 2009) available as log-ratios of paired-end *in vivo* and *in vitro* MNase data aligned to sacCer1. Missing values were replaced by the chromosome mean and values exceeding 3 times the genome-wide median were clipped to this maximal value. We smoothed the profiles using a 1D Gaussian filter with standard deviation (std) σ = 5 truncated at 3 stds. The genome-wide minimum was added to all values before scaling and discretizing the continuous values y as *x* = *floor(*2. 5 * *y*). This resulted in 22 discrete classes for both the *in vitro* and *in vivo* data, however the highest class was very rare in the *in vitro* data (observed at around 400bp) and did not feature in the training data. We obtained the Hidden-Markov model (HMM) from the same publication (Kaplan *et al*., 2009) and generated its predictions for both sacCer1 and sacCer3. We annotated the unique nucleosome map in *S.cerevisiae* from chemical cleavage sequencing of (Brogaard *et al*., 2012) to the yeast genome sacCer3(Chereji *et al*., 2018). Nucleosome dyad positions derived from ATAC-seq samples were downloaded from (Schep *et al*., 2015). We obtained raw MNase-seq data from (Henikoff *et al*., 2011) and mapped these to the sacCer3 genome with Bowtie2 (parameters: `-X 2000`), version 2.3.4.3 (Langmead and Salzberg, 2012). Resulting alignments were filtered to remove unmapped, multi-mapping and unpaired reads and duplicated fragments were removed. Next, we computed separate signal tracks for TF fragments (max length <= 80) and mono nucleosome fragments (140 <= length <= 200). Both signaling tracks were scaled to fragments per million (FPM) where the scaling factor was computed as 10^6^ divided by the number of fragments of the TF and mononuc fragments separately. Finally, signal track computation was performed using deepTools2’s “bamCoverage” function (parameters: `-bs 1 --extendReads --center`) (Ramírez *et al*., 2016). Given the significant difference in sequencing depth of the two replicates (~191 million reads vs ~31 million) we only used the deeper sequenced first replicate for all subsequent analysis. Instead of normalising the read coverage by taking the log ratio between coverage and genomic mean, we are modelling here directly the number of reads to preserve the sparsity of the TF-fragment signal. Missing values were replaced by the chromosome mean, read counts exceeding 3 times the genome-wide median were clipped to this maximal value and normalised by subtracting the genome-wide mean after which negative values were set to zero. We then smoothed the two profiles separately using a 1D Gaussian filter with σ = 5 truncated at 3 stds. We discretized these continuous values y as *x* = *floor(Z* * *y*) where Z=20 and Z=10 for the transcription factor and nucleosome profiles, respectively. This resulted in 5 and 9 discrete classes for the transcription factor and nucleosome profiles respectively.

Publicly available preprocessed high-resolution binding sites for Abf1, Cbf1, Rap1 and Reb1 as measured by ChIP-exo were downloaded from (Rossi, Lai and Pugh, 2018).

##### Human data sets

We obtained gene and TSS annotations as well as the reference sequences in hg38 from UCSC via the R Package TxDb.Hsapiens.UCSC.hg38.knownGene (Team BC, Maintainer BP, 2019). We downloaded nucleosome occupancy data from several lymphoblastoid cell lines from (Gaffney *et al*., 2012) preprocessed and aligned to the hg38 reference genome as bigWig file (in Reads Per Million, RPM) from (Zhao *et al*., 2018)(sample *hsNuc0340101*). The read coverage per base-pair around human TSSs was computed with the function “computeMatrix” in the deepTools2 package (Ramírez *et al*., 2016) (parameters: `reference-point --referencePoint TSS -a 1000 -b 1000 --binSize 1 --sortRegions keep –nanAfterEnd --missingDataAsZero`). Read counts exceeding 3 times the median of the whole dataset were clipped to this maximal value and subsequently normalised by taking the log2 ratio between read counts and genome-wide mean. We then smoothed the profiles with a 1D Gaussian filter std σ = 5 truncated at 3 stds and discretized the continuous signal y as *x* = *floo*(2. 0 * *y*). This resulted in 17 discrete classes.

Publicly available CTCF binding sites for the cell line GM12878 were downloaded from the ENCODE portal (Davis *et al*., 2018; Zhang *et al*., 2020).

Previously determined DNAse hypersensitivity quantitative trait loci (dhsQTL) were downloaded from the supplementary information of (Degner *et al*., 2012). Downloaded dhsQTLs were overlapped with promoter regions and we only retained those dsQTLs that were overlapping promoter regions (+/−1kb around TSSs).

#### Model architecture, implementation and training

##### Binary classification CNN to model nucleosome specificity

Nucleosomal DNA-sequences of 201bp were derived from the set of unique nucleosome dyads as measured by chemical cleavage (Brogaard *et al*., 2012) by extending 100bp either side from the dyad position. Dyads that were closer than 100bp to chromosome ends were excluded. This set of nucleosome sequences was then divided into training (50%), validation (30%) and test (20%) set. Reverse complements of the sequences were added to each of these datasets and then one-hot encoded. One-hot encoded means in this context that for example adenine is encoded as (1, 0, 0, 0), cytosine as (0, 1, 0, 0) and so forth. We generated a set of ‘negative’ examples of the same size by randomly shuffling each of the input sequences preserving the distribution of dinucleotides. One hot encoded nucleosomal and background sequences were used as input to train convolutional neural networks (CNN) implemented to predict labels y=1 (nucleosomal sequence) and y=0 (shuffled sequence) respectively.

CNNs were implemented with the python package Keras2.0 (Chollet and Others, 2015) with tensorflow backend. We used dropout and early stopping (patience = 20). We trained each CNN for 100 epochs with the Adam optimizer and reduced the learning rate by a factor of 0.1 if the loss had not improved for 10 epochs. In addition to the loss function, we monitored the accuracy as area under the receiver-operator curve (auROC) and Matthew’s correlation coefficient (MCC). To determine the optimal architecture of a CNN capturing sequence features of nucleosome dyad preferences *in vivo,* we searched the hyperparameter search space in Supplementary Table 1 to determine the optimal set of hyperparameters using the hyperparameter optimization algorithm “Tree of Parzen Estimators” (TPE) (Bergstra *et al*., 2011) as implemented in hyperopt package (Bergstra, Yamins and Cox, 2013). We performed 51 independent hyperparameter searches. In each of the hyperparameter searches, we performed 30 trials (e.g. iterations) and returned the model maximising the MCC on the validation dataset across those trials. As the final model, we chose the model which maximised this score across all 51 searches.

##### ChromWave models

We divided the yeast genome into windows of fixed length which formed as one-hot encoded DNA sequences the input to the ChromWave models. We split the chromosomes randomly into training, validation and test sets (excluding the mitochondrial chromosome): chromosomes II and V of sacCer1 (chromosomes V and VI of sacCer3) were used for test, chromosomes I, VII and XVI of sacCer1 (chromosomes I, VIII and XVI of sacCer3) for validation and all other chromosomes of sacCer1 (of sacCer3) used for training for nucleosome (TF-nucleosome) models. As prediction target for each sequence, the respective discretised *in vitro*, *in vivo* nucleosome occupancy profile or the TF-nucleosome profiles were used. For each sequence, the reverse complement together with the reversed occupancy profile(s) were added to the respective data set. Sequence and target pairs thus allocated to training, validation and test set were randomly shuffled within each set.

At each base-pair position the model is asked to perform a classification task to predict the class of the discretised nucleosome occupancy at that position. To combat the bias towards the overrepresented genomic mean, we use a weighted multi-class cross-entropy as loss function where the class weights were computed as follows: the class weight of class *i* was computed as the median of all class frequencies divided by the frequency of class *i.* Here, the class frequency was computed as the total number of class occurrences divided by the fragment length. Finally, to avoid very large weights for rarely occurring classes the minimum between the class weight and 100 was chosen. In the case of the TF-nucleosome model, the class weights were independently computed for each of the two profiles. Finally, we mapped the predicted classes back to the respective binned read counts in the original continuous ranges by applying the inverse transformation which was used to discretise the data (see above) on the predictions. For example, for predictions of the *in vitro* nucleosomes, the class label is multiplied by 2.5 to recover the binned read count before the signal is smoothed with a 1D Gaussian filter with σ = 5 truncated at 3 stds. Code implementing the models and hyperparameter search can be found in the ChromWave repository on github (https://github.com/luslab/ChromWave/). Models were implemented using the python package Keras 2.3.1 (Chollet and Others, 2015) with tensorflow backend.

In addition to the loss function, we monitored the class (or categorical) accuracy and Pearson’s correlation coefficient between observed and predicted profiles (after mapping class label predictions back to the continuous space as described above) on the validation set. To determine the optimal architecture of a ChromWave model, we searched the hyperparameter search spaces in Supplementary Table 1 to determine the optimal sets of hyperparameters of the model maximising the sum of class accuracy and Pearson’s correlation coefficient between observed and mapped predicted profiles in the validation set. The hyperparameter optimisation algorithm TPE (Bergstra *et al*., 2011) was used as implemented in the hyperopt package (Bergstra, Yamins and Cox, 2013). In each of the hyperparameter searches, we performed 100 trials where we trained each model for 50 epochs and then trained 6 additional neural networks on the full training set using this set of hyperparameters for 100 epochs.

For the human promoter model, we used one-hot encoded DNA-sequences of the human genome (*hg38*) corresponding to +/−1000bp around annotated TSSs as input to the model. We split the chromosomes randomly into training, test and validation data (excluding the sex and mitochondrial chromosomes). As prediction target for each sequence, the respective discretised nucleosome occupancy profile was used. For each sequence, the reverse complement together with the reversed occupancy profile was added to the respective data set. Sequence and target pairs allocated to training, validation and test set were randomly shuffled within each set. The class frequency was computed as the total number of class occurrences divided by the fragment length. Finally, to avoid very large weights for rarely occurring classes the minimum between the class weight and 40 was chosen. We employed the same training strategy as for the yeast ChromWave models. The search spaces for the hyperparameters and optimal values for each of the models are shown in Supplementary Table 1.

##### Transfer learning

In addition to training the weights of the neural networks from a random initialisation, we also compared different transfer learning approaches. These were evaluated as part of the hyperparameter search which selected the best combination of hyperparameters and transfer learning strategy.

For the *in vivo* nucleosome occupancy, we trained (i) a network was initialised with the weights of the *in vitro* nucleosome ChromWave (keeping the best architecture constant) and was trained for a further 100 epochs, (ii) networks during a hyperparameter search with (a) a set of convolutional layers are initialized with the weights of the filters of the *in vitro* nucleosome ChromWave model and subsequently frozen (ie. not further trained), or (b) trained further, and (c) a set of convolutional layers are initialized with the weights of the filters of CNNs trained on protein-binding-microarray data of 89 yeast transcription factors (Zhu *et al*., 2009) in addition to the nucleosome *in vitro* nucleosome ChromWave model filters, as described in (Alipanahi *et al*., 2015) - here all filters were further trained.

For the TF-nucleosome competition model, we compared (i) initialising all convolutional layers at random, (ii) adding the pre-trained convolutional layers of the nucleosome specificity model trained on the chemical cleavage data (but not allowing these to be trained further), and (iii) initialising a set of trainable convolutional layers with position weight matrices (and their reverse complement) of a set of yeast transcription factors.

The hyperparameter search identified the following transfer learning approaches for the training of the final models. For the *in vitro* nucleosome occupancy we initialise the first set of non-trainable convolutional layers of the ChromWave model with the weights from the convolutional layers of the CNN trained on the chemical cleavage data. For the *in vivo* nucleosome occupancy, we initialised a set of convolutional layers with the weights of the filters of the *in vitro* nucleosome ChromWave model which were not trained further. For the TF-nucleosome competition model, we added the convolutional layers of the CNN trained on the chemical cleavage data (but not allowing these to be trained further), and initialised a set of trainable convolutional layers with position weight matrices (and their reverse complement) of a set of yeast transcription factors. As set of transcription factors, we chose the following previously described nucleosome-deplacing factors: Abf1, Cbf1, McM1, Rap1, Reb1, Orc1, Asg1, Azf1, Bas1, Ecm22, Ino4, Leu3, Rfx1, Rgm1, Rgt1, Rsc3, Sfp1, Stb4, Stb5, Stp1, Sum1, Sut1, Tbf1, Tbs1, Tea1, Uga3, Ume6, Urc2 (Groups 1&2 in (Yan, Chen and Bai, 2018)). PFMs and convolutional layers of the pretrained CNNs are also available on github (https://github.com/luslab/ChromWave/).

For the human promoter nucleosome model, we trained networks during a hyperparameter search from a random initialisation as well as with a set of convolutional layers initialized with the weights of the filters of an intermediate *in vivo* nucleosome occupancy trained on the yeast data (which was not used as the final yeast model). The final human promoter model trained with the described transfer learning approach as identified by the hyperparameter search.

### QUANTIFICATION AND STATISTICAL ANALYSIS

#### Correlations between observed and predicted signals genome-wide

We computed the Pearson’s correlation coefficients in sliding windows of 300bp (with step size 1) between observed and predicted profiles. To evaluate the performance of the TF-Nucleosome competition model we computed the geometric means between the TF and nucleosomes profiles as previously published in (Zhong, Wasson and Hartemink, 2014): if *r_TF_* and *r_NUC_* are the Pearson correlation coefficients of predicted and observed TF and nucleosome signals in a 300bp window, we assign the score 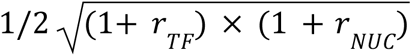 to the first base in that window. Note that this score ranges from 0 to 1. In case of the nucleosome-only data, we report genome-wide measures (e.g Pearson’s correlation coefficient) using all regions omitting bases with missing values in the MNase-seq data. This approach is not used for the TF-nucleosome model where the TF signal is very sparse and excluding regions with zero reads would highly influence this correlation and at the same time not penalise for predicted signal in unmapped regions.

#### Comparisons of occupancy-enriched and occupancy-depleted regions

To compare the predictions of the nucleosome ChromWave models with the predictions of the HMM model from (Kaplan *et al*., 2009) as well as with both the *in vitro* and *in vivo* data on the scale of individual nucleosomes, we followed a similar approach to (Kaplan *et al*., 2009): Using sliding windows of 50bp, we defined nucleosome-enriched regions as those windows whose mean occupancy is above some threshold, t_e_, and nucleosome-depleted regions as those windows whose maximum occupancy is below some threshold, t_d_. We are using the thresholds identified as optimal for the HMM model from (Kaplan *et al*., 2009) to predict *in vitro* nucleosome positions: t_e_=mean(x)+0.5*sd(x) and t_d_=mean(x)-0.5*sd(x) where x is the predicted signal genome-wide. Predictions were mapped back to the original (continuous) ranges before calculating enrichment and depletion.

In this way we defined 19,860 and 20,461 nucleosomes from the *in vivo* and *in vitro* MNase-seq data, respectively, as well as 19,577 and 20,188 from the *in vivo* and *in vitro* ChromWave predictions. To compare linker lengths, nucleosome width and signal amplitudes in mapped regions, we restricted ourselves to nucleosomes in the prediction and data profiles that overlapped by at least 100bp. This resulted in a comparison of 11,667 and 10,794 *in vivo* nucleosomes from the MNase-Seq data and ChromWave’s prediction respectively, and 13,007 12,758 *in vitro* nucleosomes from the MNase-Seq data and ChromWave’s prediction, respectively.

For TF and nucleosome occupancy, the distributions are bimodal with many regions covered by no (or very low numbers of) reads while bound regions are covered by many. Using sliding windows of 10bp and 50bp for TFs and nucleosomes respectively, we defined enriched windows if the mean occupancy score exceeded the 85th percentiles in both predictions and MNase-Seq data. Predictions were mapped back to the original (continuous) ranges before calculating enrichment. This resulted in 50,944 TF binding-sites and 48,592 nucleosome-enriched regions in the MNase-Seq data, as well as 41,432 TF binding-sites and 51,949 nucleosome enriched-regions derived from ChromWave’s predictions.

In case of the human promoter nucleosome occupancy, we defined enriched and depleted regions in the same way as in yeast (taking sliding windows of 50bp and using above/below the mean 0.5*std as threshold). In the case of the data we observed a peak of the widths of the enriched regions around 1000bp and therefore filtered out any enriched or depleted regions with more than 500bp for further analysis. This resulted in 4,290 and 54,101 nucleosome positions from the MNase-Seq data and ChromWave’s prediction, respectively.

We report several genome-wide measures to evaluate the ability of our models to distinguish between bound and unbound regions: while the genome-wide Pearson Correlation coefficient between occupancy predictions observed profiles captures the general trends of the predicted profiles, the AUCs between predicted and observed TF and nucleosome-enriched regions measure the accuracy of the model to derive the positions of TF binding-sites and nucleosomes accurately.

#### Motif filter visualisations and annotations

The first convolutional layer of a CNN can be summarised as a conventional position frequency matrix (PFM) and visualised with a seqlogo plot as previously described (Alipanahi *et al*., 2015; Angermueller *et al*., 2017). We matched these PFMs against available yeast motif databases (Zhu and Zhang, 1999; MacIsaac *et al*., 2006; Pachkov *et al*., 2013; Hume *et al*., 2015; Fornes *et al*., 2020) with TOMTOM (Gupta *et al*., 2007). Motif filters were defined as ‘known’ if they matched against a known motif with q-value < 0.05.

#### PCA embeddings of learned patterns/motifs

As a measure of importance of the learned motifs, we computed the maximum output value (maximal activation) of the respective ChromWave filter for each DNA-sequence window. To understand the impact of these motifs on ChromWave’s TF predictions better, we computed the correlation between the vector of filter activations along the sequence and the predicted TF occupancy profile. A positive correlation implies the existence of the motif in a sequence is increasing TF binding, while a negative correlation implies the motif being unfavourable for TF binding. We then applied PCA to the maximal activations of all filters across all DNA-sequence windows.

#### Clustering of learned patterns/motifs across models

Motif clustering was performed as previously described in (Vierstra *et al*., 2020) with code obtained from https://github.com/jvierstra/motif-clustering. Briefly, motif similarities were computed between filters from all yeast models (*in vitro, in vivo, TF-NUC,* excluding the pre-trained filters of each model) using TOMTOM (Gupta *et al*., 2007). These pairwise similarities were used as input for hierarchical clustering followed by cluster definition by cutting the dendrogram on a given height (h=0.9). These clusters were manually annotated using previous TOMTOM results and visualized. Results can be found in Supplementary Table 2.

#### *In silico* mutagenesis scores and visualisation

To visualise the changes in prediction given changes in input, we computed the prediction profile for each of the possible base changes at each position along the input sequence as previously described (Alipanahi *et al*., 2015). To visualise the effect of each base change in an area of interest, we summed over all changes within that area of interest for each base. We visualize the result as a heatmap where each column is a position in the sequence, each row a substituted base pair and the colour reflecting this summed change across the sequence if that base pair is substituted in the sequence. To visualise global changes as line plots, we computed the maximum (‘gain’) and minimum (‘loss’) difference of the prediction given of all possible changes at each position.

#### DHS accessibility under *in silico* mutagenesis

We used the human-promoter trained ChromWave model to elucidate the impact of different SNPs in previously annotated *DNase I sensitivity quantitative trait loci* (dsQTLs) (Degner *et al*., 2012). Here we substituted each possible base-change in the position of the dsQTL and computed the difference in prediction on the dsQTL of the model. To account for overall-changes to the binding profile we also computed the Pearson’s correlation coefficient between the predicted WT profiles and the prediction given the perturbed input sequence. dsQTLs were differentiated into “within DHS” and “outside DHS” depending on their position relative to the DHS which is influenced by the respective SNP (defined in the same publication (Degner *et al*., 2012)). Results of this analysis can be found in Supplementary Table 3.

#### Other visualisations

Heatmaps and average meta-profiles were produced using the “computeMatrix” and subsequent “plotHeatmap” or “plotProfile” functions in the in the deepTools2 package (Ramírez *et al*., 2016).

## Notes

https://github.com/luslab/ChromWave

https://github.com/luslab/chromWaveR

### Competing Interest Statement

The authors have declared no competing interest.

### Summary of Updates

General clarifications and small corrections throughout the text; Typo in Fig 2A corrected.

## References

Alipanahi, B. et al. (2015) ‘Predicting the sequence specificities of DNA- and RNA-binding proteins by deep learning’, Nature biotechnology, 33(8), pp. 831–838.

Almouzni, G. and Wolffe, A.P. (1995) ‘Constraints on transcriptional activator function contribute to transcriptional quiescence during early Xenopus embryogenesis’, The EMBO journal, 14(8), pp. 1752–1765.

Angermueller, C. et al. (2017) ‘DeepCpG: accurate prediction of single-cell DNA methylation states using deep learning’, Genome biology, 18(1), p. 67.

Avsec, Ž., Weilert, M., et al. (2021) ‘Base-resolution models of transcription-factor binding reveal soft motif syntax’, Nature genetics, 53(3), pp. 354–366.

Avsec, Ž., Agarwal, V., et al. (2021) ‘Effective gene expression prediction from sequence by integrating long-range interactions’, Nature methods, 18(10), pp. 1196–1203.

Awazu, A. (2017) ‘Prediction of nucleosome positioning by the incorporation of frequencies and distributions of three different nucleotide segment lengths into a general pseudo k-tuple nucleotide composition’, Bioinformatics, 33(1), pp. 42–48.

Badis, G. et al. (2008) ‘A library of yeast transcription factor motifs reveals a widespread function for Rsc3 in targeting nucleosome exclusion at promoters’, Molecular cell, 32(6), pp. 878–887.

Bartke, T. et al. (2010) ‘Nucleosome-interacting proteins regulated by DNA and histone methylation’, Cell, 143(3), pp. 470–484.

Bergstra, J.S. et al. (2011) ‘Algorithms for Hyper-Parameter Optimization’, in J. Shawe-Taylor et al. (eds) Advances in Neural Information Processing Systems 24. Curran Associates, Inc., pp. 2546–2554.

Bergstra, J., Yamins, D. and Cox, D. (2013) ‘Making a Science of Model Search: Hyperparameter Optimization in Hundreds of Dimensions for Vision Architectures’, in International Conference on Machine Learning. International Conference on Machine Learning, pp. 115–123.

Brogaard, K. et al. (2012) ‘A map of nucleosome positions in yeast at base-pair resolution’, Nature, 486(7404), pp. 496–501.

Charoensawan, V. et al. (2012) ‘DNA sequence preferences of transcriptional activators correlate more strongly than repressors with nucleosomes’, Molecular cell, 47(2), pp. 183–192.

Chereji, R.V. et al. (2018) ‘Precise genome-wide mapping of single nucleosomes and linkers in vivo’, Genome biology, 19(1), p. 19.

Chollet, F. and Others (2015) *Keras*. Available at: https://keras.io.

Davis, C.A. et al. (2018) ‘The Encyclopedia of DNA elements (ENCODE): data portal update’, Nucleic acids research, 46(D1), pp. D794–D801.

Degner, J.F. et al. (2012) ‘DNase I sensitivity QTLs are a major determinant of human expression variation’, Nature, 482(7385), pp. 390–394.

Durinck, S. et al. (2005) ‘BioMart and Bioconductor: a powerful link between biological databases and microarray data analysis’, Bioinformatics, 21(16), pp. 3439–3440.

Durinck, S. et al. (2009) ‘Mapping identifiers for the integration of genomic datasets with the R/Bioconductor package biomaRt’, Nature protocols, 4(8), pp. 1184–1191.

Engel, S.R. et al. (2014) ‘The reference genome sequence of Saccharomyces cerevisiae: then and now’, G3, 4(3), pp. 389–398.

Ernst, J. and Kellis, M. (2017) ‘Chromatin-state discovery and genome annotation with ChromHMM’, Nature protocols, 12(12), pp. 2478–2492.

Floer, M. et al. (2010) ‘A RSC/nucleosome complex determines chromatin architecture and facilitates activator binding’, Cell, 141(3), pp. 407–418.

Fornes, O. et al. (2020) ‘JASPAR 2020: update of the open-access database of transcription factor binding profiles’, Nucleic acids research, 48(D1), pp. D87–D92.

Gaffney, D.J. et al. (2012) ‘Controls of nucleosome positioning in the human genome’, PLoS genetics, 8(11), p. e1003036.

Ganapathi, M. et al. (2011) ‘Extensive role of the general regulatory factors, Abf1 and Rap1, in determining genome-wide chromatin structure in budding yeast’, Nucleic acids research, 39(6), pp. 2032–2044.

Gkikopoulos, T. et al. (2011) ‘A role for Snf2-related nucleosome-spacing enzymes in genome-wide nucleosome organization’, Science, 333(6050), pp. 1758–1760.

Gupta, S. et al. (2007) ‘Quantifying similarity between motifs’, Genome biology, 8(2), p. R24.

Gupta, S. et al. (2008) ‘Predicting human nucleosome occupancy from primary sequence’, PLoS computational biology, 4(8), p. e1000134.

Hartley, P.D. and Madhani, H.D. (2009) ‘Mechanisms that specify promoter nucleosome location and identity’, Cell, 137(3), pp. 445–458.

Hayes, J.J. and Wolffe, A.P. (1992) ‘The interaction of transcription factors with nucleosomal DNA’, BioEssays: news and reviews in molecular, cellular and developmental biology, 14(9), pp. 597–603.

He, K. et al. (2016) ‘Deep Residual Learning for Image Recognition’, in 2016 IEEE Conference on Computer Vision and Pattern Recognition (CVPR), pp. 770–778.

Henikoff, J.G. et al. (2011) ‘Epigenome characterization at single base-pair resolution’, Proceedings of the National Academy of Sciences of the United States of America, 108(45), pp. 18318–18323.

He, X. et al. (2013) ‘Contribution of nucleosome binding preferences and co-occurring DNA sequences to transcription factor binding’, BMC genomics, 14, p. 428.

Hume, M.A. et al. (2015) ‘UniPROBE, update 2015: new tools and content for the online database of protein-binding microarray data on protein-DNA interactions’, Nucleic acids research, 43(Database issue), pp. D117–22.

Jayaram, N., Usvyat, D. and R Martin, A.C. (2016) ‘Evaluating tools for transcription factor binding site prediction’, BMC bioinformatics, 17(1), p. 547.

John, S. et al. (2008) ‘Interaction of the Glucocorticoid Receptor with the Chromatin Landscape’, Molecular Cell, pp. 611–624. Available at: https://doi.org/10.1016/j.molcel.2008.02.010.

John, S. et al. (2011) ‘Chromatin accessibility pre-determines glucocorticoid receptor binding patterns’, Nature genetics, 43(3), pp. 264–268.

Kaplan, N. et al. (2009) ‘The DNA-encoded nucleosome organization of a eukaryotic genome’, Nature, 458(7236), pp. 362–366.

Kaplan, T. et al. (2011) ‘Quantitative models of the mechanisms that control genome-wide patterns of transcription factor binding during early Drosophila development’, PLoS genetics, 7(2), p. e1001290.

Kasprzyk, A. et al. (2004) ‘EnsMart: a generic system for fast and flexible access to biological data’, Genome research, 14(1), pp. 160–169.

Kelley, D.R. et al. (2018) ‘Sequential regulatory activity prediction across chromosomes with convolutional neural networks’, Genome research, 28(5), pp. 739–750.

Kelley, D.R., Snoek, J. and Rinn, J. (2015) ‘Basset: Learning the regulatory code of the accessible genome with deep convolutional neural networks’, bioRxiv. Available at: https://doi.org/10.1101/028399.

Kent, N.A. et al. (1994) ‘Chromatin structure modulation in Saccharomyces cerevisiae by centromere and promoter factor 1’, Molecular and cellular biology, 14(8), pp. 5229–5241.

Kim, H.D. and O’Shea, E.K. (2008) ‘A quantitative model of transcription factor-activated gene expression’, Nature structural & molecular biology, 15(11), pp. 1192–1198.

Kim, T.H. et al. (2007) ‘Analysis of the vertebrate insulator protein CTCF-binding sites in the human genome’, Cell, 128(6), pp. 1231–1245.

Krietenstein, N. et al. (2016) ‘Genomic Nucleosome Organization Reconstituted with Pure Proteins’, Cell, 167(3), pp. 709–721.e12.

Lam, F.H., Steger, D.J. and O’Shea, E.K. (2008) ‘Chromatin decouples promoter threshold from dynamic range’, Nature, 453(7192), pp. 246–250.

Langmead, B. and Salzberg, S.L. (2012) ‘Fast gapped-read alignment with Bowtie 2’, Nature methods, 9(4), pp. 357–359.

Liu, G. et al. (2016) ‘A deformation energy-based model for predicting nucleosome dyads and occupancy’, Scientific reports, 6, p. 24133.

MacIsaac, K.D. et al. (2006) ‘An improved map of conserved regulatory sites for Saccharomyces cerevisiae’, BMC bioinformatics, 7, p. 113.

Mathelier, A. and Wasserman, W.W. (2013) ‘The next generation of transcription factor binding site prediction’, PLoS computational biology, 9(9), p. e1003214.

Mirny, L. (2009) ‘Nucleosome-mediated cooperativity between transcription factors’, Nature Precedings [Preprint]. Available at: https://doi.org/10.1038/npre.2009.2796.1.

Neph, S. et al. (2012) ‘An expansive human regulatory lexicon encoded in transcription factor footprints’, Nature, 489(7414), pp. 83–90.

van den Oord, A., Kalchbrenner, N., et al. (2016) ‘Conditional Image Generation with PixelCNN Decoders’, *arXiv [cs.CV]*. Available at: http://arxiv.org/abs/1606.05328.

van den Oord, A., Dieleman, S., et al. (2016) ‘WaveNet: A Generative Model for Raw Audio’, arXiv [cs.SD]. Available at: http://arxiv.org/abs/1609.03499.

Ozonov, E.A. and van Nimwegen, E. (2013) ‘Nucleosome free regions in yeast promoters result from competitive binding of transcription factors that interact with chromatin modifiers’, PLoS computational biology, 9(8), p. e1003181.

Pachkov, M. et al. (2013) ‘SwissRegulon, a database of genome-wide annotations of regulatory sites: recent updates’, Nucleic acids research, 41(Database issue), pp. D214–20.

Peckham, H.E. et al. (2007) ‘Nucleosome positioning signals in genomic DNA’, Genome research, 17(8), pp. 1170–1177.

Quang, D. and Xie, X. (2016) ‘DanQ: a hybrid convolutional and recurrent deep neural network for quantifying the function of DNA sequences’, Nucleic acids research, 44(11), p. e107.

Ramírez, F. et al. (2016) ‘deepTools2: a next generation web server for deep-sequencing data analysis’, Nucleic acids research, 44(W1), pp. W160–5.

Raveh-Sadka, T. et al. (2012) ‘Manipulating nucleosome disfavoring sequences allows fine-tune regulation of gene expression in yeast’, Nature genetics, 44(7), pp. 743–750.

Rossi, M.J., Lai, W.K.M. and Pugh, B.F. (2018) ‘Genome-wide determinants of sequence-specific DNA binding of general regulatory factors’, Genome research, 28(4), pp. 497–508.

Schep, A.N. et al. (2015) ‘Structured nucleosome fingerprints enable high-resolution mapping of chromatin architecture within regulatory regions’, Genome research, 25(11), pp. 1757–1770.

Schübeler, D. (2015) ‘Function and information content of DNA methylation’, Nature, 517(7534), pp. 321–326.

Schwessinger, R. et al. (2020) ‘DeepC: predicting 3D genome folding using megabase-scale transfer learning’, Nature methods, 17(11), pp. 1118–1124.

Segal, E. et al. (2006) ‘A genomic code for nucleosome positioning’, Nature, 442(7104), pp. 772–778.

Segal, E. and Widom, J. (2009) ‘Poly(dA:dT) tracts: major determinants of nucleosome organization’, Current opinion in structural biology, 19(1), pp. 65–71.

Smedley, D. et al. (2015) ‘The BioMart community portal: an innovative alternative to large, centralized data repositories’, Nucleic acids research, 43(W1), pp. W589–98.

Stewart, A.J., Hannenhalli, S. and Plotkin, J.B. (2012) ‘Why transcription factor binding sites are ten nucleotides long’, Genetics, 192(3), pp. 973–985.

Svaren, J. et al. (1994) ‘Analysis of the competition between nucleosome formation and transcription factor binding’, The Journal of biological chemistry, 269(12), pp. 9335–9344.

Team BC, Maintainer BP (2019) ‘TxDb.Hsapiens.UCSC.hg38.knownGene: Annotation package for TxDb object(s)’, *R package version 3.4.6.* [Preprint]. Available at: https://bioconductor.org/packages/release/data/annotation/html/TxDb.Hsapiens.UCSC.hg38.knownGene.html.

Tsankov, A. et al. (2011) ‘Evolutionary divergence of intrinsic and trans-regulated nucleosome positioning sequences reveals plastic rules for chromatin organization’, Genome research, 21(11), pp. 1851–1862.

Tsankov, A.M. et al. (2010) ‘The role of nucleosome positioning in the evolution of gene regulation’, PLoS biology, 8(7), p. e1000414.

Vierstra, J. et al. (2020) ‘Global reference mapping of human transcription factor footprints’, Nature, 583(7818), pp. 729–736.

Wang, M. et al. (2018) ‘DeFine: deep convolutional neural networks accurately quantify intensities of transcription factor-DNA binding and facilitate evaluation of functional non-coding variants’, Nucleic acids research, 46(11), p. e69.

Wasserman, W.W. and Sandelin, A. (2004) ‘Applied bioinformatics for the identification of regulatory elements’, Nature reviews. Genetics, 5(4), pp. 276–287.

Wasson, T. and Hartemink, A.J. (2009) ‘An ensemble model of competitive multi-factor binding of the genome’, Genome research, 19(11), pp. 2101–2112.

Yan, C., Chen, H. and Bai, L. (2018) ‘Systematic Study of Nucleosome-Displacing Factors in Budding Yeast’, Molecular cell, 71(2), pp. 294–305.e4.

Yarragudi, A. et al. (2004) ‘Comparison of ABF1 and RAP1 in chromatin opening and transactivator potentiation in the budding yeast Saccharomyces cerevisiae’, Molecular and cellular biology, 24(20), pp. 9152–9164.

Zeng, H. and Gifford, D.K. (2017) ‘Predicting the impact of non-coding variants on DNA methylation’, Nucleic acids research [Preprint]. Available at: https://doi.org/10.1093/nar/gkx177.

Zerbino, D.R. et al. (2018) ‘Ensembl 2018’, Nucleic Acids Research, pp. D754–D761. Available at: https://doi.org/10.1093/nar/gkx1098.

Zhang, J. et al. (2020) ‘An integrative ENCODE resource for cancer genomics’, Nature communications, 11(1), p. 3696.

Zhang, Y. et al. (2009) ‘Intrinsic histone-DNA interactions are not the major determinant of nucleosome positions in vivo’, Nature structural & molecular biology, 16(8), pp. 847–852.

Zhang, Z. et al. (2011) ‘A packing mechanism for nucleosome organization reconstituted across a eukaryotic genome’, Science, 332(6032), pp. 977–980.

Zhao, Y. et al. (2018) ‘NucMap: a database of genome-wide nucleosome positioning map across species’, Nucleic acids research [Preprint]. Available at: https://doi.org/10.1093/nar/gky980

Zhong, J., Wasson, T. and Hartemink, A.J. (2014) ‘Learning protein-DNA interaction landscapes by integrating experimental data through computational models’, Bioinformatics, 30(20), pp. 2868–2874.

Zhu, C. et al. (2009) ‘High-resolution DNA-binding specificity analysis of yeast transcription factors’, Genome research, 19(4), pp. 556–566.

Zhu, F. et al. (2018) ‘The interaction landscape between transcription factors and the nucleosome’, Nature [Preprint]. Available at: https://doi.org/10.1038/s41586-018-0549-5.

Zhu, J. and Zhang, M.Q. (1999) ‘SCPD: a promoter database of the yeast Saccharomyces cerevisiae’, Bioinformatics, 15(7-8), pp. 607–611.

