## Supplementary Tables for "ChromWave: Deciphering the DNA-encoded competition between transcription factors and nucleosomes with deep neural networks": KEY RESOURCES TABLE.docx

| REAGENT or RESOURCE | SOURCE | IDENTIFIER |
| --- | --- | --- |
| Deposited Data |  |  |
| Nucleosome dyads (Chemical Cleavage) (*S. cerevisia*e) | (Brogaard et al., 2012) | https://www.ncbi.nlm.nih.gov/pmc/articles/PMC3786739/bin/NIHMS370046-supplement-4.txt |
| Nucleosome dyads (Chemical Cleavage) (*S. cerevisia*e) | (Chereji et al., 2018) | [https://www.ncbi.nlm.nih.gov/geo/query/acc.cgi?acc=GSE97290](https://media.nature.com/original/nature-assets/nature/journal/v486/n7404/extref/nature11142-s2.txt) |
| ATAC-Seq (*S. cerevisia*e) | (Schep et al., 2015) | [https://www.ncbi.nlm.nih.gov/geo/download/?acc=GSE66386&format=file&file=GSE66386%5Fall%5Fnucpos%2Ebed%2Etar%2Egz](https://media.nature.com/original/nature-assets/nature/journal/v486/n7404/extref/nature11142-s2.txt) |
| MNase-Seq (*S. cerevisia*e) | (Henikoff et al., 2011) | <https://www.ncbi.nlm.nih.gov/geo/query/acc.cgi?acc=GSE30551> |
| MNase-Seq (*S. cerevisia*e) | (Kaplan et al., 2009) | [https://genie.weizmann.ac.il/pubs/nucleosomes08/nucleosomes08_data.html](https://media.nature.com/original/nature-assets/nature/journal/v486/n7404/extref/nature11142-s2.txt) |
| MNase-Seq (*human*) | (Gaffney et al., 2012; Zhao et al., 2018) | [https://bigd.big.ac.cn/nucmap/NucMap_FTP_Directory/Homo_sapiens/byDataType/Transformed_read_signals/Homo_sapiens.hsNuc0340101.nucleosome.shift.bw](https://media.nature.com/original/nature-assets/nature/journal/v486/n7404/extref/nature11142-s2.txt) |
| dsQTLs | (Degner et al., 2012) | [http://eqtl.uchicago.edu/dsQTL_data/](https://media.nature.com/original/nature-assets/nature/journal/v486/n7404/extref/nature11142-s2.txt) |
| DNA-hypersensitive sites | (Degner et al., 2012) | [https://www.ncbi.nlm.nih.gov/geo/download/?acc=GSE31388&format=file&file=GSE31388%5FdsQtlTable%2Etxt%2Egz](https://www.google.com/url?q=https://www.ncbi.nlm.nih.gov/geo/download/?acc%3DGSE31388%26format%3Dfile%26file%3DGSE31388%255FdsQtlTable%252Etxt%252Egz&sa=D&ust=1610996533378000&usg=AOvVaw3278Y4EJEQcj4uUO7YU1Ei) |
| Chip-exo (*S. cerevisia*e) | (Rossi et al., 2018) | [ftp://ftp.ncbi.nlm.nih.gov/geo/series/GSE93nnn/GSE93662/suppl/](https://media.nature.com/original/nature-assets/nature/journal/v486/n7404/extref/nature11142-s2.txt) |
| ChIP-Seq (human) | (Davis et al., 2018) | [https://www.encodeproject.org/files/ENCFF960ZGP/@@download/ENCFF960ZGP.bed.gz](https://media.nature.com/original/nature-assets/nature/journal/v486/n7404/extref/nature11142-s2.txt) |
| Protein-binding microarray (*S. cerevisia*e) | (Zhu et al., 2009) | [http://thebrain.bwh.harvard.edu/uniprobe/downloads/GR09/GR09_deBruijn.zip](https://media.nature.com/original/nature-assets/nature/journal/v486/n7404/extref/nature11142-s2.txt) |
| Experimental Models: Organisms/Strains | | |
| sacCer1 (SGD reference R27.1.1) | (Engel et al., 2014) | <http://hgdownload.cse.ucsc.edu/goldenpath/sacCer1/bigZips/> |
| sacCer3 (SGD reference R64.2.1) | (Engel et al., 2014) | <http://hgdownload.cse.ucsc.edu/goldenpath/sacCer3/bigZips/> |
| hg38 | (Team BC, Maintainer BP) | <https://bioconductor.org/packages/release/data/annotation/html/TxDb.Hsapiens.UCSC.hg38.knownGene.html> |
| Software and Algorithms | | |
| Code repository | This manuscript | <https://github.com/luslab/ChromWave> |
| Code repository | This manuscript | <https://github.com/luslab/chromWaveR> |
| DeepTools2 | (Ramírez et al., 2016) | <https://deeptools.readthedocs.io/en/develop/> |
| Keras | (Chollet and Others, 2015) | <https://keras.io/> |
| Hyperopt | (Bergstra et al., 2013) | <http://hyperopt.github.io/hyperopt/> |
| biomaRt | (Durinck et al., 2005, 2009) | <https://www.bioconductor.org/packages/release/bioc/html/biomaRt.html> |
| Bowtie2 | (Langmead and Salzberg, 2012) | <http://bowtie-bio.sourceforge.net/bowtie2/index.shtml> |
| R package TxDb.Hsapiens.UCSC.hg38.knownGene | (Team BC, Maintainer BP, 2019) | <https://bioconductor.org/packages/release/data/annotation/html/TxDb.Hsapiens.UCSC.hg38.knownGene.html> |
| TOMTOM | (Gupta et al., 2007) | <http://meme-suite.org/tools/tomtom> |
| Rtracklayer | (Lawrence et al., 2009) | <https://bioconductor.org/packages/release/bioc/html/rtracklayer.html> |
| Other | | |

Bergstra, J., Yamins, D., and Cox, D. (2013). Making a Science of Model Search: Hyperparameter Optimization in Hundreds of Dimensions for Vision Architectures. In International Conference on Machine Learning, pp. 115–123.

Brogaard, K., Xi, L., Wang, J.-P., and Widom, J. (2012). A map of nucleosome positions in yeast at base-pair resolution. Nature *486*, 496–501.

Chereji, R.V., Ramachandran, S., Bryson, T.D., and Henikoff, S. (2018). Precise genome-wide mapping of single nucleosomes and linkers in vivo. Genome Biol. *19*, 19.

Chollet, F., and Others (2015). Keras.

Davis, C.A., Hitz, B.C., Sloan, C.A., Chan, E.T., Davidson, J.M., Gabdank, I., Hilton, J.A., Jain, K., Baymuradov, U.K., Narayanan, A.K., et al. (2018). The Encyclopedia of DNA elements (ENCODE): data portal update. Nucleic Acids Res. *46*, D794–D801.

Degner, J.F., Pai, A.A., Pique-Regi, R., Veyrieras, J.-B., Gaffney, D.J., Pickrell, J.K., De Leon, S., Michelini, K., Lewellen, N., Crawford, G.E., et al. (2012). DNase I sensitivity QTLs are a major determinant of human expression variation. Nature *482*, 390–394.

Durinck, S., Moreau, Y., Kasprzyk, A., Davis, S., De Moor, B., Brazma, A., and Huber, W. (2005). BioMart and Bioconductor: a powerful link between biological databases and microarray data analysis. Bioinformatics *21*, 3439–3440.

Durinck, S., Spellman, P.T., Birney, E., and Huber, W. (2009). Mapping identifiers for the integration of genomic datasets with the R/Bioconductor package biomaRt. Nat. Protoc. *4*, 1184–1191.

Engel, S.R., Dietrich, F.S., Fisk, D.G., Binkley, G., Balakrishnan, R., Costanzo, M.C., Dwight, S.S., Hitz, B.C., Karra, K., Nash, R.S., et al. (2014). The reference genome sequence of Saccharomyces cerevisiae: then and now. G3 *4*, 389–398.

Gaffney, D.J., McVicker, G., Pai, A.A., Fondufe-Mittendorf, Y.N., Lewellen, N., Michelini, K., Widom, J., Gilad, Y., and Pritchard, J.K. (2012). Controls of nucleosome positioning in the human genome. PLoS Genet. *8*, e1003036.

Gupta, S., Stamatoyannopoulos, J.A., Bailey, T.L., and Noble, W.S. (2007). Quantifying similarity between motifs. Genome Biol. *8*, R24.

Henikoff, J.G., Belsky, J.A., Krassovsky, K., MacAlpine, D.M., and Henikoff, S. (2011). Epigenome characterization at single base-pair resolution. Proc. Natl. Acad. Sci. U. S. A. *108*, 18318–18323.

Kaplan, N., Moore, I.K., Fondufe-Mittendorf, Y., Gossett, A.J., Tillo, D., Field, Y., LeProust, E.M., Hughes, T.R., Lieb, J.D., Widom, J., et al. (2009). The DNA-encoded nucleosome organization of a eukaryotic genome. Nature *458*, 362–366.

Langmead, B., and Salzberg, S.L. (2012). Fast gapped-read alignment with Bowtie 2. Nat. Methods *9*, 357–359.

Lawrence, M., Gentleman, R., and Carey, V. (2009). rtracklayer: an R package for interfacing with genome browsers. Bioinformatics *25*, 1841–1842.

Ramírez, F., Ryan, D.P., Grüning, B., Bhardwaj, V., Kilpert, F., Richter, A.S., Heyne, S., Dündar, F., and Manke, T. (2016). deepTools2: a next generation web server for deep-sequencing data analysis. Nucleic Acids Res. *44*, W160–W165.

Rossi, M.J., Lai, W.K.M., and Pugh, B.F. (2018). Genome-wide determinants of sequence-specific DNA binding of general regulatory factors. Genome Res. *28*, 497–508.

Schep, A.N., Buenrostro, J.D., Denny, S.K., Schwartz, K., Sherlock, G., and Greenleaf, W.J. (2015). Structured nucleosome fingerprints enable high-resolution mapping of chromatin architecture within regulatory regions. Genome Res. *25*, 1757–1770.

Team BC, Maintainer BP (2019). TxDb.Hsapiens.UCSC.hg38.knownGene: Annotation package for TxDb object(s). R Package Version 3.4.6.

Team BC, Maintainer BP TxDb.Hsapiens.UCSC.hg38.knownGene: Annotation package for TxDb object(s).

Zhao, Y., Wang, J., Liang, F., Liu, Y., Wang, Q., Zhang, H., Jiang, M., Zhang, Z., Zhao, W., Bao, Y., et al. (2018). NucMap: a database of genome-wide nucleosome positioning map across species. Nucleic Acids Res.

Zhu, C., Byers, K.J.R.P., McCord, R.P., Shi, Z., Berger, M.F., Newburger, D.E., Saulrieta, K., Smith, Z., Shah, M.V., Radhakrishnan, M., et al. (2009). High-resolution DNA-binding specificity analysis of yeast transcription factors. Genome Res. *19*, 556–566.
